# Evidence of microglial involvement in the childhood abuse-associated increase in perineuronal nets in the ventromedial prefrontal cortex

**DOI:** 10.1101/2024.03.08.584135

**Authors:** Claudia Belliveau, Reza Rahimian, Gohar Fakhfouri, Clémentine Hosdey, Sophie Simard, Maria Antonietta Davoli, Dominique Mirault, Bruno Giros, Gustavo Turecki, Naguib Mechawar

**Author notes:** Corresponding Author: Naguib Mechawar, PhD, Douglas Mental Health University Institute, Department of Psychiatry, McGill University. shared first authorship (Claudia Belliveau, Reza Rahimian).

## Abstract

Microglia, known for their diverse roles in the central nervous system, have recently been recognized for their involvement in degrading the extracellular matrix. Perineuronal nets (PNNs), a specialized form of this matrix, are crucial for stabilizing neuronal connections and constraining plasticity. Our group recently reported increased PNN densities in the ventromedial prefrontal cortex (vmPFC) of depressed individuals that died by suicide in adulthood after experiencing childhood abuse (DS-CA) compared to matched controls. To explore potential underlying mechanisms, we employed a comprehensive approach in similar postmortem vmPFC samples, combining a human matrix metalloproteinase and chemokine array, isolation of CD11b-positive microglia and enzyme-linked immunosorbent assays (ELISA). Our findings indicate a significant downregulation of matrix metalloproteinase (MMP)-9 and tissue inhibitors of metalloproteinases (TIMP)-2 in both whole vmPFC grey matter and isolated microglial cells from DS-CA samples. Furthermore, our experiments reveal that a history of child abuse is associated with diminished levels of microglial CX3CR1 and IL33R in both vmPFC whole lysate and CD11b isolated cells. However, levels of the CX3CR1 ligand, CX3CL1 (Fractalkine), did not differ between groups. While these data suggest potential long-lasting alterations in microglial markers in the vmPFC of individuals exposed to severe childhood adversity, direct functional assessments were not conducted. Nonetheless, these findings offer insight into how childhood abuse may contribute to PNN alterations via microglial-related mechanisms.

**Graphical Abstract:** 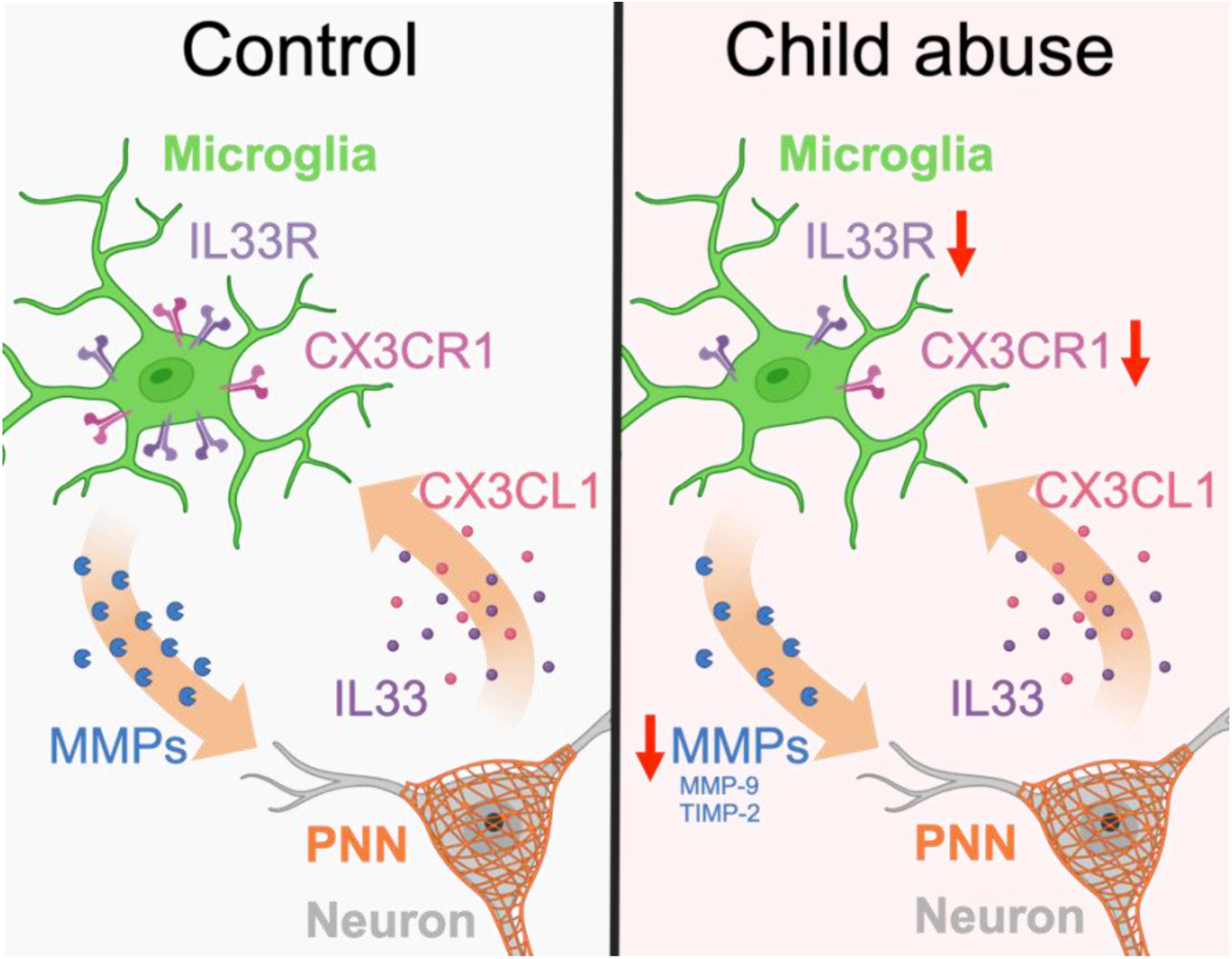

## 1. Introduction

Microglia, the brain’s resident immune cells, are heterogeneous and continuously surveying their environment, responding to inflammatory insults or changes in brain states. Rather than being passively “activated”, microglia engage in dynamic interactions that can contribute to the modulation of neuroinflammation (Javanmehr et al., 2022; Paolicelli et al., 2022; Wright-Jin and Gutmann, 2019). A growing body of evidence suggests that aberrant neuroinflammation may play an instrumental role in major depressive disorder (MDD) (Li et al., 2022a; Rahimian et al., 2022; Rahimian et al., 2021; Troubat et al., 2021), at least in certain patients, through the chronic priming of microglia in limbic regions (Torres-Platas et al., 2014). Microglial development can also be affected by early-life stress (ELS) after which these cells can exhibit altered phagocytosis, morphology and homeostatic processing (Bolton et al., 2022; Catale et al., 2020; Rahimian et al., 2024a; Reemst et al., 2022). Furthermore, microglial cells respond to environmental signals such as chronic stress by releasing various molecules (cytokines, chemokines and growth factors) (Schramm and Waisman, 2022). Microglial cells also regulate synaptic pruning in an activity dependent manner engulfing presynaptic inputs which is crucial for proper brain development and function (Schafer et al., 2012). Microglia have recently gained attention for their role in regulating brain plasticity through modulation of the extracellular matrix (ECM) (reviewed in (Crapser et al., 2021; Dzyubenko and Hermann, 2023)). One specialized form of condensed ECM, perineuronal nets (PNNs), function to inhibit axon growth and spine formation (Monnier et al., 2003; Pearson et al., 2018) while acting as a scaffolding for neuronal circuitry (Sigal et al., 2019; Tsien, 2013). Emerging evidence implicates microglia as key players in ECM degradation, with a strong focus on PNNs (Arreola et al., 2021; Crapser et al., 2021; Crapser et al., 2020a; Crapser et al., 2020b; Nguyen et al., 2020; Rahimian et al., 2024a; Sun et al., 2023; Venturino et al., 2021). The mechanisms underlying this phenomenon, however, remain incompletely understood. Instances of direct PNN regulation through microglial phagocytosis have been recognized, as evidenced by PNN staining (*Wisteria Floribunda Lectin* (WFL) or anti-aggrecan) visible in CD68+ lysosomes of microglia in healthy brain (Nguyen et al., 2020; Sun et al., 2023; Venturino et al., 2021) and spinal cord (Tansley et al., 2022), Alzheimer’s Disease (Crapser et al., 2020b), Huntington’s Disease (Crapser et al., 2020a) and adult-onset leukoencephalopathy with axonal spheroids and pigmented glia (Arreola et al., 2021) brains. The existing body of literature highlights the importance of interactions between neurons and microglial cells in the intricate regulation of PNN structures. The interaction between neuronal IL33 and microglial IL33-receptor (IL33R) is a good example. This interaction has been shown to trigger microglial engulfment of PNN component aggrecan; an effect abolished in knockout mice of either the neuronal ligand or microglial receptor (Nguyen et al., 2020). Tansley et al. investigated the role of microglia in peripheral nerve injury-mediated PNN degradation. They demonstrated that PNN engulfment similarly occurs through neuronal CX3CL1 binding to microglial CX3CR1. Knocking out either CX3CL1 or CX3CR1 prevents PNN degradation and consequent accumulation of WFL in CD68+ microglial lysosomes (Tansley et al., 2022), highlighting that microglia-neuron interactions play a role in microglial engulfment of PNNs.

Matrix metalloproteinases (MMPs) are enzymes that catalyze glycoproteins in the ECM. Their activity is tightly controlled in the healthy brain by endogenous tissue inhibitors (TIMP-1, -2, -3, -4) that bind to their catalytic site (Laronha and Caldeira, 2020). MMPs can be classified, based on their affinity to various substrates, into different subgroups: gelatinases (MMP-2, -9), stromelysins (MMP-3, -10 and -11) and collagenases (MMP-1, -8, -13, -18) among others (Laronha and Caldeira, 2020). Importantly, all classes of MMPs have already been implicated in ECM regulation. Specifically, they have all been found to target aggrecan core protein, tenascins and link proteins which are all fundamental components of PNNs (Könnecke and Bechmann, 2013; Laronha and Caldeira, 2020). Different types of MMPs, especially MMP-2 and MMP-9, are expressed by microglial cells and their roles have previously been established in neuroinflammatory cascades (Könnecke and Bechmann, 2013). The most studied MMP, MMP-9, plays a fundamental role in neurodevelopment (myelination) and homeostasis in adulthood (Bitanihirwe and Woo, 2020). Studies show that MMP-9, released by glia and neurons, is directly responsible for PNN degradation, as shown by increased densities of PNNs following genetic deletion of the MMP-9 gene in mice (Wen et al., 2017). Interestingly, antidepressant efficacy in stressed mice has also been linked to PNN degradation by MMP-9 (Alaiyed et al., 2019).

In a recent postmortem study, we identified in depressed suicides a child abuse-associated increase in WFL^+^ PNNs in the lower layers of the ventromedial prefrontal cortex (vmPFC) (Tanti et al., 2022) a cortical region critically involved in decision-making, impulsivity and suicidal tendencies (Hiser and Koenigs, 2018; Teicher et al., 2016). The current study sought to explore the impact of child abuse on the regulation of PNNs by microglia in the vmPFC. Using an unbiased MMP antibody array the relative expression of various MMPs in vmPFC was quantified using both grey matter lysate and CD11b-positive (CD11b^+^) isolated microglia/macrophages. Additionally, microglial markers implicated in ECM modulation such as CX3CR1 and IL33R were quantified using enzyme-linked immunosorbent assay (ELISA) in both types of samples. We hypothesized that the expression of microglial proteins involved in ECM degradation would be downregulated in depressed suicides with a history of child abuse because of the increased PNN density we observed previously.

## 2. Methods

### 2.1 Human brain samples

The postmortem human brain samples used in this study were provided by the Douglas-Bell Canada Brain Bank (DBCBB) (Montreal, Canada) with ethical approval from the Douglas Research Ethics Board (IUSMD-20-35). Brains, donated by familial consent, were acquired by the DBCBB thanks to a collaboration between the McGill Group for Suicide Studies and the Quebec Coroner’s Office. A panel of psychiatrists created clinical vignettes for each individual by compiling all available information obtained from medical records, the coroner’s report, toxicological analyses, and other sources of information, as described previously (A. Dumais et al., 2005). Samples used in this study were from psychiatrically healthy controls (CTRL) and depressed suicides with a history of severe child abuse (DS-CA). Non-random childhood abuse (sexual, physical and neglect) was assessed through a standardized psychological autopsy with next of kin, using a modified version of the Childhood Experience of Care and Abuse (CECA) (Bifulco et al., 1994). Only the most severe cases (score of 1-2) occurring before the age of 15 were included in the DS-CA group. Any indication of neurodevelopmental, neurological, or other co-morbid mental illnesses were cause for exclusion.

Brains were cut into 0.5cm-thick coronal sections upon arrival at the DBCBB then flash-frozen in isopentane at -35°C and stored at -80°C until use, referred to hereafter as archived frozen brain samples. Expert staff at the DBCBB dissected vmPFC (Brodmann area 11/12) at the equivalent of plate 3 (approximately -48 mm from the center of the anterior commissure) of the Mai and Paxinos Atlas of the Human Brain (Juergen Mai, 2016). Group characteristics of archived frozen brain samples are outlined in **Table 1**.

**Table 1:**
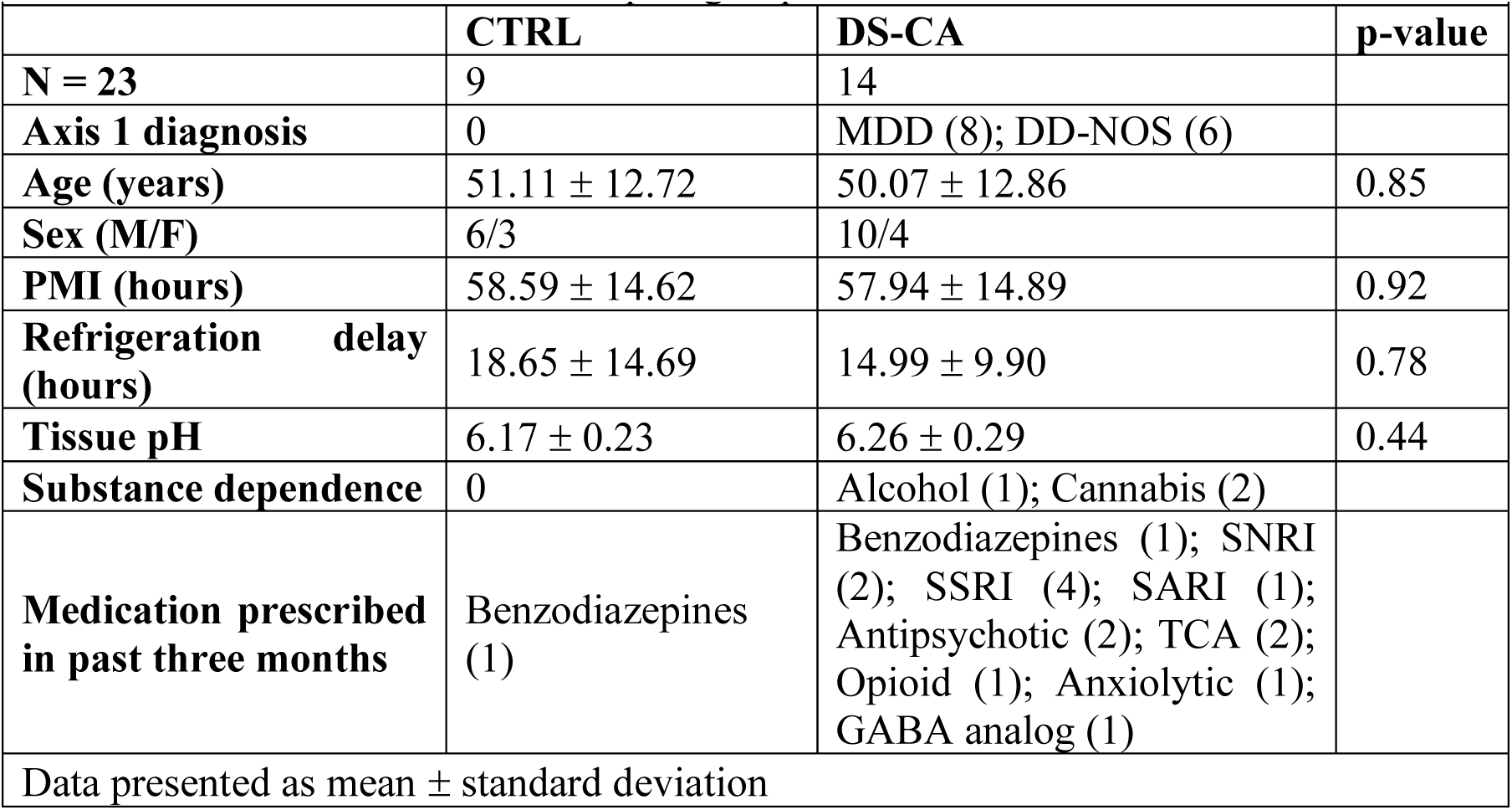
Archived frozen brain samples group characteristics.

| **Table 1:** Archived frozen brain samples group characteristics |  |  |  |
| --- | --- | --- | --- |
|  | **CTRL** | **DS-CA** | **p-value** |
| **N = 23** | 9 | 14 |  |
| **Axis 1 diagnosis** | 0 | MDD (8); DD-NOS (6) |  |
| **Age (years)** | 51.11 ± 12.72 | 50.07 ± 12.86 | 0.85 |
| **Sex (M/F)** | 6/3 | 10/4 |  |
| **PMI (hours)** | 58.59 ± 14.62 | 57.94 ± 14.89 | 0.92 |
| **Refrigeration delay (hours)** | 18.65 ± 14.69 | 14.99 ± 9.90 | 0.78 |
| **Tissue pH** | 6.17 ± 0.23 | 6.26 ± 0.29 | 0.44 |
| **Substance dependence** | 0 | Alcohol (1); Cannabis (2) |  |
| **Medication prescribed in past three months** | Benzodiazepines (1) | Benzodiazepines (1); SNRI (2); SSRI (4); SARI (1); Antipsychotic (2); TCA (2); Opioid (1); Anxiolytic (1); GABA analog (1) |  |
| Data presented as mean ± standard deviation |  |  |  |

### 2.2 Human MMP Antibody Array

Grey matter was dissected from archived frozen vmPFC samples and lysed using RIPA buffer (Sigma, St. Louis, MO, USA) and then protein concentration was measured on a TECAN microplate reader. A human MMP antibody array (RayBiotech® C-Series, Human MMP Array C1, Norcross, GA, USA) was used to compare the relative abundance of 10 MMPs and their inhibitors (TIMPs) between CTRL and DS-CA samples according to the manufacturer’s protocol. This array allows for the semi-quantitative analysis of MMP-1, -2, -3, -8, -9, -10 and -13 as well as of TIMP-1, -2 and -4. Unfortunately, MMP-10 was removed from the analyses because quantities were too low and outside of the quantifiable range. It is noteworthy that the RayBiotech® human MMP array detects both pro and active forms of MMP and our findings cannot discriminate between active and non-active forms. However, this array was the only technique available that provided enough sensitivity, robustness and consistency to detect multiple MMPs in post-mortem human brain samples. An antibody array membrane was used per subject, each processed in separate wells of the provided incubation tray, printed side up. Membranes were incubated under constant agitation for 30 mins at room temperature (RT) with 2 ml of blocking buffer. Next, 1 ml of diluted sample containing 150μg of protein was added to each well and incubated overnight at 4°C under constant agitation. Membranes were washed for 5 minutes at RT thrice with Wash Buffer 1 and twice with Wash Buffer 2. Membranes were then incubated with 1 ml of the prepared Biotinylated Antibody Cocktail and incubated overnight at 4°C under constant agitation. Wash Buffer 1 + 2 were used as described above. Membranes were incubated for 2h at RT with 1X HRP-Streptavidin followed by Wash Buffer 1 + 2. Lastly, chemiluminescence detection was completed using the Detection Buffers provided. 2-D densitometry was conducted using ImageJ to detect the spot signal densities. Relative expression was calculated by subtracting background from each spot and normalizing to a reference positive control spot. Each antibody was spotted in duplicate on the membrane so normalized relative expression was averaged per subject to obtain a single measurement per antibody.

### 2.3 Human Chemokine Antibody Array

Grey matter was dissected from archived frozen vmPFC samples and lysed using RIPA buffer (Sigma, St. Louis, MO, USA) and then protein concentration was measured on a TECAN microplate reader. A human chemokine antibody array (RayBiotech®, Human Chemokine Array C1, Norcross, GA, USA) was used to compare the relative abundance of CX3CL1 and several important chemokines including CCL2, CCL3, CCL4 and CCL5 between CTRL and DS-CA samples according to the manufacturer’s protocol. The workflow of this array is the same as the MMP array which was explained in the previous section.

### 2.4 ELISA

ELISA was employed for quantitative measurement of human CD68 (cluster of differentiation 68), CX3CR1 (CX3C motif chemokine receptor 1), Cathepsin-S (Cat-S), TREM2 (triggering receptor expressed on myeloid cells 2), and IL33R (interleukin 33 receptor/ST2) in vmPFC samples following the manufacturer’s instructions. In brief, grey matter was dissected from the archived frozen vmPFC and lysed using RIPA buffer (Sigma, St. Louis, MO, USA). Total protein concentration for each sample was measured using a BCA Protein Assay kit (cat # 23225, Thermo Fisher Scientific, Saint-Laurent, Quebec, Canada). Depending on the assay, 50-100 μg total protein and standards were added to antibody pre-coated wells and after addition of conjugated antibodies specific to the analyte of interest (sandwich ELISA) or conjugated-analyte in the case of CD68 (competitive ELISA) microplates were incubated at 37°C for 60 minutes. Substrate addition led to color development in proportion to the amount of the analyte, except for CD68 kit, where the intensity of the color (optical density, OD) was inversely proportional to the CD68 concentration. The reaction was terminated by addition of acidic stop solution and absorbance was read at 450 nm. A standard curve was plotted relating the OD to the concentration of standards. The analyte concentration (pg/ml or ng/ml) in each sample was then interpolated from the corresponding standard curve. Optimal sample protein amounts for each ELISA assay were determined in a pilot experiment to ensure OD values would fall within the range of the corresponding standard curve. A detailed description of the procedure is found on MyBioSource.com.

### 2.5 Western Blotting

The same samples were used for ELISA and Western Blotting. Pilot tests were conducted to find out whether chondroitinase ABC (chABC) digestion of the samples was necessary prior to immunoblotting. We observed no discernible differences between samples subjected to various durations of chABC digestion (20h, 1h, 30mins) and those left untreated, and thus omitted this step to streamline the process and optimize time efficiency. 4-20% Mini-PROTEAN® TGX Stain-Free Protein, 12 well, 20 μl gels (cat # 4568095, Bio-Rad, Hercules, CA, USA) were used to separate proteins based on size. Blots were probed with an antibody for the N-terminal (DIPEN) neo-epitope of aggrecan core protein cleaved by MMP between amino acids PEN341 and 342FFG (clone BC-4, 1:100, cat # 1042002, MDBioproducts, Oakdale, MN, USA). Prior to gel loading, samples were prepared by adding 1X Laemmli Buffer and incubated at 95°C for 5 minutes, followed by centrifugation and subsequent placement on ice for 3 minutes. Equal volumes were loaded into the gradient gels (each sample 35 μg of protein) and electrophoresis was conducted for 50 minutes at 135 mV in a Bio-Rad Mini-PROTEAN® Tetra Cell electrophoresis chamber containing 1X running buffer, with constant agitation facilitated by a magnetic stir bar. Preceding the 30-minute semi-dry standard protein transfer from the gel to a nitrocellulose membrane, stain-free gels were activated for 45 seconds on a Chemi-doc to capture a total protein control image. Following a 1-hour blocking step in 5% milk in PBS-tween, the primary antibody, also diluted in the same solution, was incubated with the blot overnight at 4°C under continuous agitation. Post-incubation, three 10-minute washes in PBS-Tween were performed, and a secondary Amersham ECL sheep anti-mouse, horseradish peroxidase linked IgG antibody (1:1000, cat # NA931, Cytvia, Marlborough, MA, USA) was subsequently incubated in 1% milk in PBS-tween for 2 hours at room temperature, again under constant agitation. Blots were subjected to three additional 10-minute washes in PBS-Tween before being exposed to an enhanced chemiluminescence (ECL) solution (1:1) for 1 minute. The blots were imaged on the Chemi-doc system using predefined settings. Original images of the blots, ladder and their stain-free gel can be found in **Supplementary File 1**.

Subsequent blot analyses adhered to the guidelines provided by Bio-Rad. Each blot incorporated a reference lane comprising a composite of all samples, serving as an inter-gel control. To succinctly outline the quantification process, the adjusted volume of the 50 kDa band was used to determine the protein mean intensity on the blot, while the adjusted volume for the entire lane on the stain-free gel provided a measure of total protein. Normalization of each sample was achieved by establishing a normalization factor, computed as the ratio of the total protein content of the sample to the adjusted volume of its corresponding stain-free gel. Subsequently, this factor was divided by the corresponding value for the reference lane, yielding a fold-difference. The resultant fold-differences were then compared between the CTRL and DS-CA.

### 2.6 Fluorescent in situ hybridization (FISH) in human vmPFC samples

Frozen vmPFC blocks were cryo-sectioned in serial 10 µm-thick sections that were collected on Superfrost charged slides. *In situ* hybridization was performed using Advanced Cell Diagnostics RNAscope® probes and reagents following the manufacturer’s guidelines. Slides were removed from the -80 °C freeze and immediately fixed in cold 10% neutral buffered formalin for 15 min. Next, samples were dehydrated by increasing gradient of ethanol baths and then air-dried for 5 min. Hydrogen peroxide incubation for 10 min quenched endogenous peroxidase activity. Next, followed by a 30 min protease digestion. The following probes were then hybridized for 2 h at 40 °C in a humidity-controlled oven: Hs-P2RY12 (ACDbio, cat # 450391), Hs-CD11b (ACDbio, cat # 555091) and Hs-GAS6 (ACDbio, REF;455141). Amplifiers were then added using the proprietary AMP reagents and the signal visualized through probe-specific HRP-based detection by tyramide signal amplification (TSA, ACDbio) with Opal dyes (Opal 520 and Opal 570; Perkin Elmer) diluted 1:600. To eliminate endogenous autofluorescence from lipofuscin and cellular debris, sections were incubated with TrueBlack (Biotium, cat # 23007, Fremont, California, USA) for 30 seconds. Slides were then cover slipped with Vectashield mounting medium with 4′,6-diamidino-2-phenylindole (DAPI) for nuclear staining (Vector Laboratories, cat # H-1800, Newark, California, USA) and kept at 4 °C until imaging. Both positive and negative controls provided by the supplier (ACDbio) were used on separate adjacent sections to confirm signal specificity. The slides were imaged at 20X magnification using the Evident Scientific VS120 Slide Scanner and the scans were transferred to QuPath (v.0.3.0) for further analysis. Regions of interest within vmPFC grey matter were demarcated manually. QuPath automated cell detection was used to outline DAPI staining and then RNAscope signal quantification. For each probe, only cells bearing three or more fluorescent puncta were counted as positively labeled. We investigated the percentage of CD11b^+^ cells expressing P2RY12 in attempt to estimate what percentage of isolated CD11b^+^ cells (next section) were microglia (P2RY12^+^). Additionally, we examined the percentage of P2RY12^+^ cells co-expressing Gas6.

### 2.7 CD11b^+^ microglia isolation using magnetic beads

To measure the relative expression of different MMPs in vmPFC microglia, fresh brain tissue was acquired on a rolling basis from the DBCBB from December 2019 – July 2023. Upon arrival at the DBCBB, samples underwent immediate processing for CD11b+ microglia isolation, followed by MMP antibody array analysis and ELISA, according to the following procedure. Miltenyi Biotec CD11b MicroBeads (cat # 130-093-634, Gaithersburg, MD, USA) were first used to isolate microglia. These beads were developed for the isolation or depletion of human and mouse cells based on their CD11b expression. In humans, CD11b is strongly expressed on myeloid cells such as cerebral microglia and weakly expressed on NK cells and some activated lymphocytes. Using a 1 cm^3^ block of fresh-non-frozen vmPFC, grey matter was dissected and CD11b^+^ microglia were isolated using a combination of enzymatic and mechanical dissociations following Miltenyi Biotec instructions. The Neural Tissue Dissociation Kit (Papain, cat # 130-092-628, Miltenyi Biotec, Gaithersburg, MD, USA) and the gentleMACS™ Dissociator (cat # 130-093-235, Miltenyi Biotec, Gaithersburg, MD, USA) are effective in generating single-cell suspensions from neural tissues prior to subsequent applications, such as MACS® Cell Separation. After mechanical and enzymatic dissociation, red blood cells were eliminated using Red Blood Cell Lysis Solution (cat # 130-094-183, Miltenyi Biotec, Gaithersburg, MD, USA). These steps were followed by application of Debris Removal Solution, which is a ready-to-use density gradient reagent (cat # 130-109-398, Miltenyi Biotec, Gaithersburg, MD, USA), allowing for a fast and effective removal of cell debris from viable cells after dissociation of various tissue types, while applying full acceleration and full brake during centrifugation. Prior to the positive selection of CD11b^+^ cells, Myelin Removal Beads were employed for the specific removal of myelin debris from single-cell suspensions (cat # 130-096-433, Miltenyi Biotec, Gaithersburg, MD, USA). Then, the CD11b^+^ cells were magnetically labeled with CD11b MicroBeads and the cell suspension was loaded onto a MACS® Column, which was placed in the magnetic field of a MACS Separator. The magnetically labeled CD11b^+^ cells were retained on the column, while the unlabeled cell fraction, which was depleted of CD11b^+^ cells ran through the column. The magnetically retained CD11b^+^ cells were finally eluted following removal of the column from the magnetic field. Lastly, the CD11b^+^ cells were processed for Human MMP antibody array and ELISA (CX3CR1 and IL33R) as described above. After rigorous characterization of the brain samples by psychological autopsy, as outlined previously, four subjects met inclusion criteria for DS-CA and could be matched to four healthy CTRL samples (**Table 2**).

**Table 2:** Fresh brain samples group characteristics.

| **Table 2:** Fresh brain samples group characteristics |  |  |  |
| --- | --- | --- | --- |
|  | **CTRL** | **DS-CA** | **p-value** |
| **N = 8** | 4 | 4 |  |
| **Axis 1 diagnosis** | 0 | MDD (2); DD-NOS (2) |  |
| **Age (years)** | 53.25 ± 12.87 | 55.5 ± 6.45 | 0.77 |
| **Sex (M/F)** | 2/2 | 2/2 |  |
| **PMI (hours)** | 51.52 ± 7.37 | 49.63 ± 2.50 | 0.66 |
| **Refrigeration delay (hours)** | 11.93 ± 6.17 | 14.51 ± 3.48 | 0.50 |
| **Tissue pH** | 6.45 ± 0.33 | 5.89 ± 0.25 | 0.23 |
| **Substance dependence** | 0 | 0 |  |
| **Medication prescribed in past three months** | Benzodiazepines (1) | SSRI (2); Anxiolytic (1); Benzodiazepine (1) |  |
| Data presented as mean ± standard deviation |  |  |  |

### 2.8 Immunostaining

A fresh 1 cm^3^ histological block was also acquired alongside the sample processed for CD11b^+^ cell isolation and fixed in 10% neutral buffered formalin for 24 h at 4 °C. Fresh-fixed blocks were then transferred into 30% sucrose in PBS until the block sank. Tissue was then flash-frozen in isopentane at -35 °C and stored at -80 °C until all subject information was available. After full characterization, the same 8 subjects that were used for CD11b^+^ cell isolation were processed for immunofluorescent staining. Fresh-fixed tissue blocks were cryo-sectioned into 40 µm-thick free-floating sections stored in cryoprotectant. Free-floating sections were washed thrice in PBS and then incubated overnight at 4 °C under constant agitation with a combination of primary antibodies diluted in blocking solution of PBS/0.2% Triton-X/2% normal donkey serum. Tissues were incubated with either anti-TMEM119 (Abcam, cat # ab185333, 1:500), anti-aggrecan (ACAN, Millipore, MAB5284, 1:500) and anti-parvalbumin (Swant, cat # PVG213, 1:500) and anti-CD68 (Abcam, cat # ab201340, 1:500). Sections were then rinsed and incubated with the appropriate fluorophore-conjugated secondary antibody: Alexa-Cy3 anti-mouse (Jackson ImmunoResearch, 715-165-151, 1:500), Alexa-488 anti-goat (Jackson ImmunoResearch, 705-545-147, 1:500), Dylight-647 anti-rabbit (Jackson ImmunoResearch, 711-605-152, 1:500) in the same blocking solution for 2 h at room temperature. Sections were once again washed thrice in PBS and endogenous autofluorescence was quenched with TrueBlack (Biotium, cat # 23007, Fremont, California, USA) for 80 seconds. Lastly, sections were mounted on Superfrost charged slides and coverslipped using Vectashield Vibrance mounting media with DAPI (Vector Laboratories, cat # H-1800, Newark, California, USA).

Whole vmPFC sections were imaged on an Evident Scientific VS120 Slide Scanner at 20X. Samples stained with anti-TMEM119 and anti-CD68 were analyzed as follows. Grey matter microglia (TMEM119^+^) were marked as CD68^high^ or CD68^low^ based on anti-CD68 expression and then the percentage of each subtype was compared between groups (CTRL vs DS-CA). 1590 TMEM119^+^ microglia were analyzed in this study, evenly distributed across groups and subjects.

To explore the spatial relationships between PNNs and microglia (samples stained with anti-TMEM119, anti-ACAN and anti-PVG14), we developed a script (Python v3.8) to classify and analyze these biological structures based on their spatial coordinates. 352 PNNs across both groups were manually localized in vmPFC grey matter and manually categorized as PNNs using QuPath Point Tool. TMEM119^+^ microglia were manually localized and categorized as Microglia using QuPath Point Tool. QuPath Points were converted to QuPath Detections and their X and Y coordinates were exported. Our script segregated PNNs and Microglia into two distinct categories based on their respective X and Y coordinates, further distinguishing PNNs based on their association with PV cells (PNNPV) or other cell types (PNNother) which were also manually marked. Each structure was assigned a unique identifier to facilitate the tracking throughout the analysis.

We then calculated the Euclidean distance between each PNN and the nearest Microglia, enabling the quantification of their spatial proximity. Distances were calculated only when they were below a specific threshold (MAX_DISTANCE = 50µm), focusing the analysis on probable biologically relevant interactions. Furthermore, we examined whether microglia were closer or not to PNNs enwrapping PV cells or other cell types, we saw no difference, so the classifications were combined into one group “PNN”. The source code can be found in **Supplementary File 2**.

### 2.9 Statistical analyses

Statistical analyses were conducted using GraphPad Prism v10.1.1 (Boston, MA, USA) and SPSS v 29.0 (IBM Corp, Armonk, New York, USA). We identified outliers utilizing Grubb’s method and assessed the distribution and homogeneity of variances through Shapiro–Wilk and Levene’s tests, respectively. Independent samples t-tests were employed for all analyses with the exception being the comparison of time of death Cathepsin-S levels between groups where two-way analysis of variance (ANOVA) was applied (CTRL: n = 9, DS-CA: n = 14). Age showed no correlation with any of the presented results. pH exhibited correlations with IL33R and TREM2 levels, while the number of medications demonstrated correlations with whole grey-matter TIMP-2 levels. Additionally, the type of antidepressant and the number of medications were correlated with CX3CR1 levels (**Supplementary Table 1**). Postmortem interval (PMI) was significantly correlated with microglial MMP-2 and TIMP-2 levels (**Supplementary Table 2**). Consequently, a one-way analysis of covariance (ANCOVA) was applied, treating group as a fixed factor and pH, PMI or medications as a covariate for these variables. A two-tailed approach was used for all tests and p-values < 0.05 were considered significant. All data is presented as mean ± standard error of the mean unless otherwise specified.

## 3. Results

### 3.1 Effect of childhood adversity on MMP levels in vmPFC grey matter whole lysate

A representative blot of human MMP antibody array used to assess the relative expression of MMPs and TIMPs in vmPFC grey matter is illustrated in **Fig. 1A**. This approach allowed us to measure a significant overall downregulation of MMPs (pro- and active form) in DS-CA compared to CTRLs. Notably, gelatinase MMP-9 exhibited a significant downregulation in DS-CA (P = 0.011) whereas MMP-2 was similar between groups (P = 0.43, **Fig. 1B**). Stromelysin MMP-3 was also significantly downregulated in DS-CA vs CTRL samples (P = 0.012, **Fig. 1C**), as was collagenases MMP-1 (P = 0.036) and MMP-8 (P = 0.001, **Fig. 1D**). MMP-13 (P = 0.08) remained similar between groups (**Fig. 1D**). Interestingly, all three MMP inhibitors displayed significant downregulation in DS-CA: TIMP-1 (P = 0.005), TIMP-2 (ANCOVA: F (1,19) = 28.14, P < 0.001) and TIMP-4 (P = 0.003, **Fig. 1E**).

**Figure 1:**
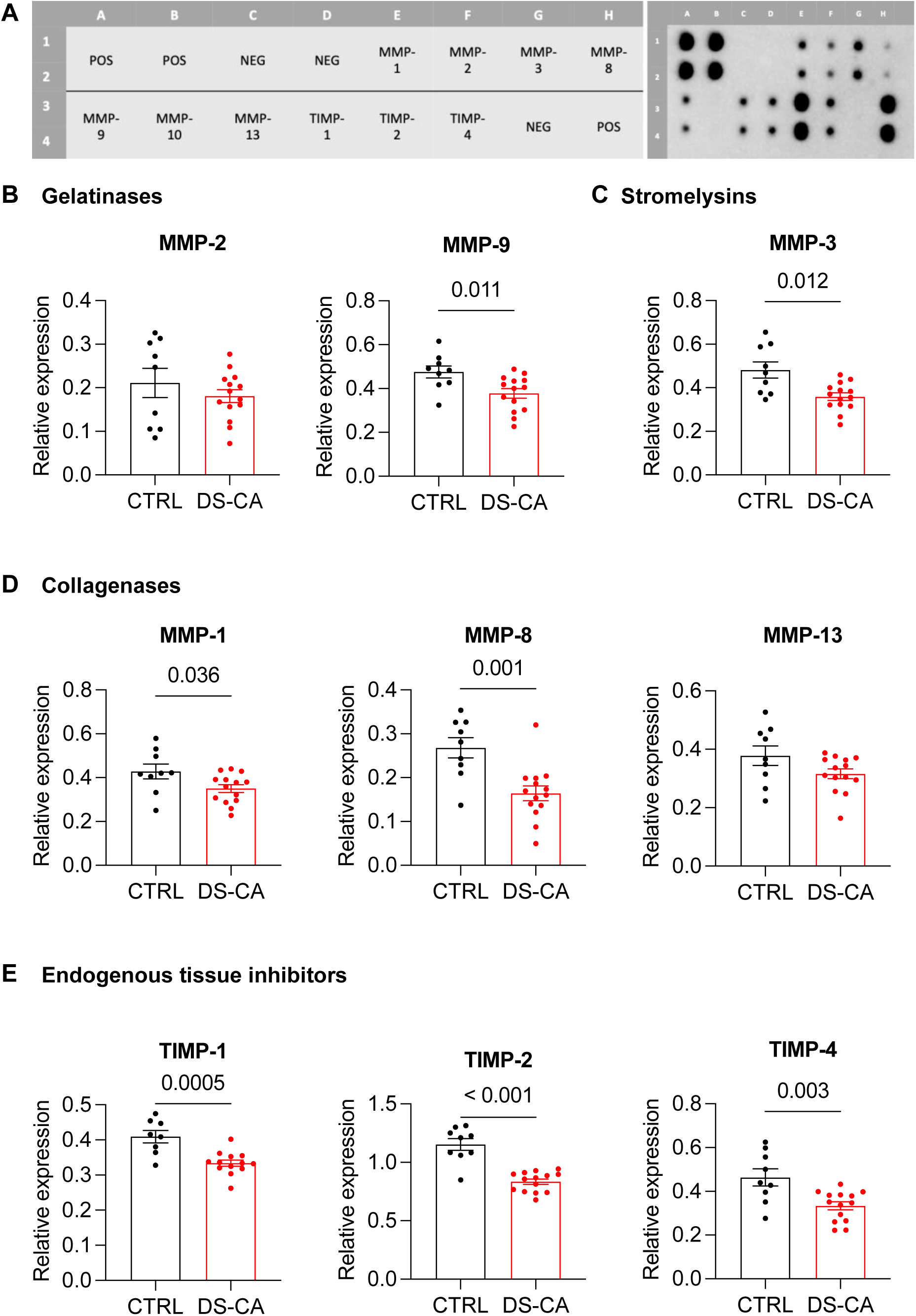
Most of the examined matrix metalloproteinases and their inhibitors are downregulated in vmPFC grey matter lysate from depressed suicides with a history of child abuse compared to matched controls. **A** Map of the human MMP Array (RayBiotec®) used and a representative blot. Each antibody is spotted in duplicate on the membrane, mean intensity was averaged per antibody as described in the Methods section. B MMP-9, C MMP-3, D MMP-1 and MMP-8 and E TIMP-1, -2, -4 are significantly decreased in DS-CA compared to CTRL. vmPFC: ventromedial prefrontal cortex, POS: positive control, NEG: negative control, MMP: matrix metalloproteinase, TIMP: endogenous tissue inhibitor, CTRL: controls, DS-CA: depressed suicides with a history of child abuse

### 3.2 Effect of childhood adversity of levels of canonical chemokines in vmPFC gray matter whole lysate

Using a similar antibody array for chemokines, we measured the relative protein expression of several interesting chemokines including CX3CL1, CCL2, CCL3, CCL4 and CCL5 (**Fig. 2A**). We were especially interested in measuring the expression of Fractalkine (CX3CL1). It is a transmembrane chemokine expressed mainly by neurons and signals through its unique receptor, CX3CR1, that is expressed in microglia (Pawelec et al., 2020) and whose signaling is involved in PNN degradation (Tansley et al., 2022). Our results revealed that there are no significance differences between the levels of CX3CL1 (ANCOVA: F(1, 19): 0.01, P = 0.93), CCL2 (P = 0.43), CCL3 (P = 0.34), CCL4 (P = 0.57) and CCL5 (ANCOVA: F(1,19) = 0.05, P = 0.83) in vmPFC lysate of DS-CA compared to CTRL (**Fig. 2B-F**).

**Figure 2:**
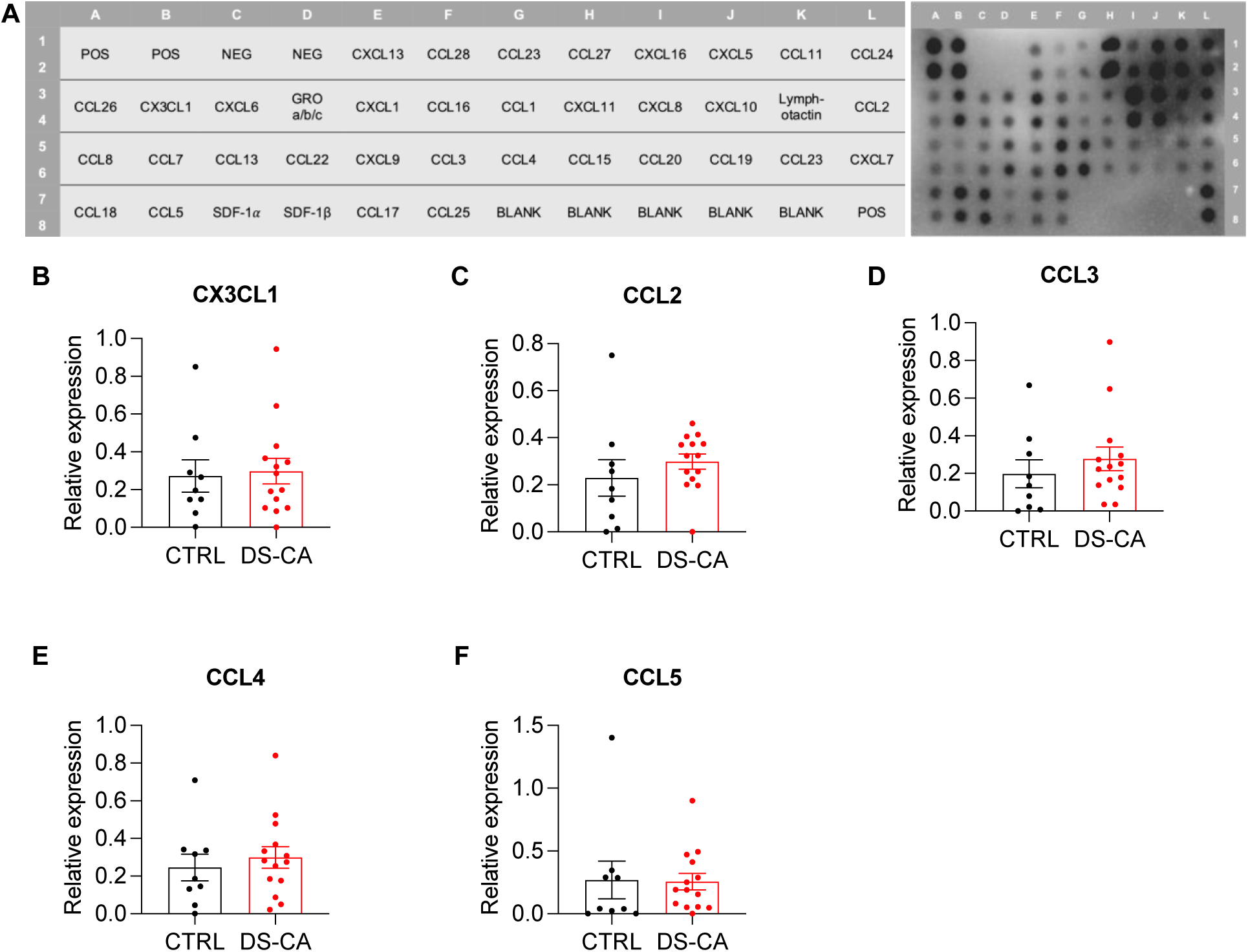
Relative protein expression of several chemokines were similar between groups in vmPFC grey matter lysate. **A** Map of the human Chemokine Array ((RayBiotec®) used and a representative blot. Each antibody is spotted in duplicate on the membrane, mean intensity was averaged per antibody as described in the Methods section. **B-F** CX3CL1, CCL2, CCL3, CCL4 and CCL5 exhibited no difference in relative expression between CTRL and DS-CA. vmPFC: ventromedial prefrontal cortex, POS: positive control, NEG: negative control, CCL: chemokine (C-C motif) ligand, CTRL: controls, DS-CA: depressed suicides with a history of child abuse

### 3.3 Effect of childhood adversity on the levels of microglia/macrophage markers involved in the regulation of PNNs in vmPFC grey matter whole lysate

We then conducted ELISAs to assess other molecules known to be involved in ECM modulation (Arreola et al., 2021; Crapser et al., 2020a; Crapser et al., 2020b; Nguyen et al., 2020; Sun et al., 2023; Tansley et al., 2022; Venturino et al., 2021). CX3CR1 (ANCOVA: F(1,19) = 45.49, P < 0.001, **Fig. 3A**) and IL33R (ANCOVA: F(1,20) = 12.81, P = 0.0020, **Fig. 3B**) were both significantly downregulated in DS-CA compared to CTRL vmPFC. Conversely, CD68 (P = 0.30, **Fig. 3C**), TREM2 (ANCOVA: F(1,20) = 0.14, P = 0.91, **Fig. 3D**) and Cat-S (P = 0.062, **Fig. 3E**) levels were similar between groups. To account for the diurnal rhythm of Cat-S (Pantazopoulos et al., 2020), subjects were classified into two categories based on time of death (6:00AM to 5:59PM: Day; 6:00PM-5:59AM: Night). Although there was no significant interaction between group and time of death for Cat-S concentration, a general reduction in Cat-S levels was observed in DS-CA subjects. Notably, individuals who died at night exhibited significantly lower Cat-S concentration (time of death x group F(1,19) = 1.53, P = 0.23; time of death F(1,19) = 2.40, P = 0.14; group F(1,19) = 5.55, P = 0.029; followed by Fisher’s LSD post-hoc test, **Fig. 3F**).

**Figure 3:**
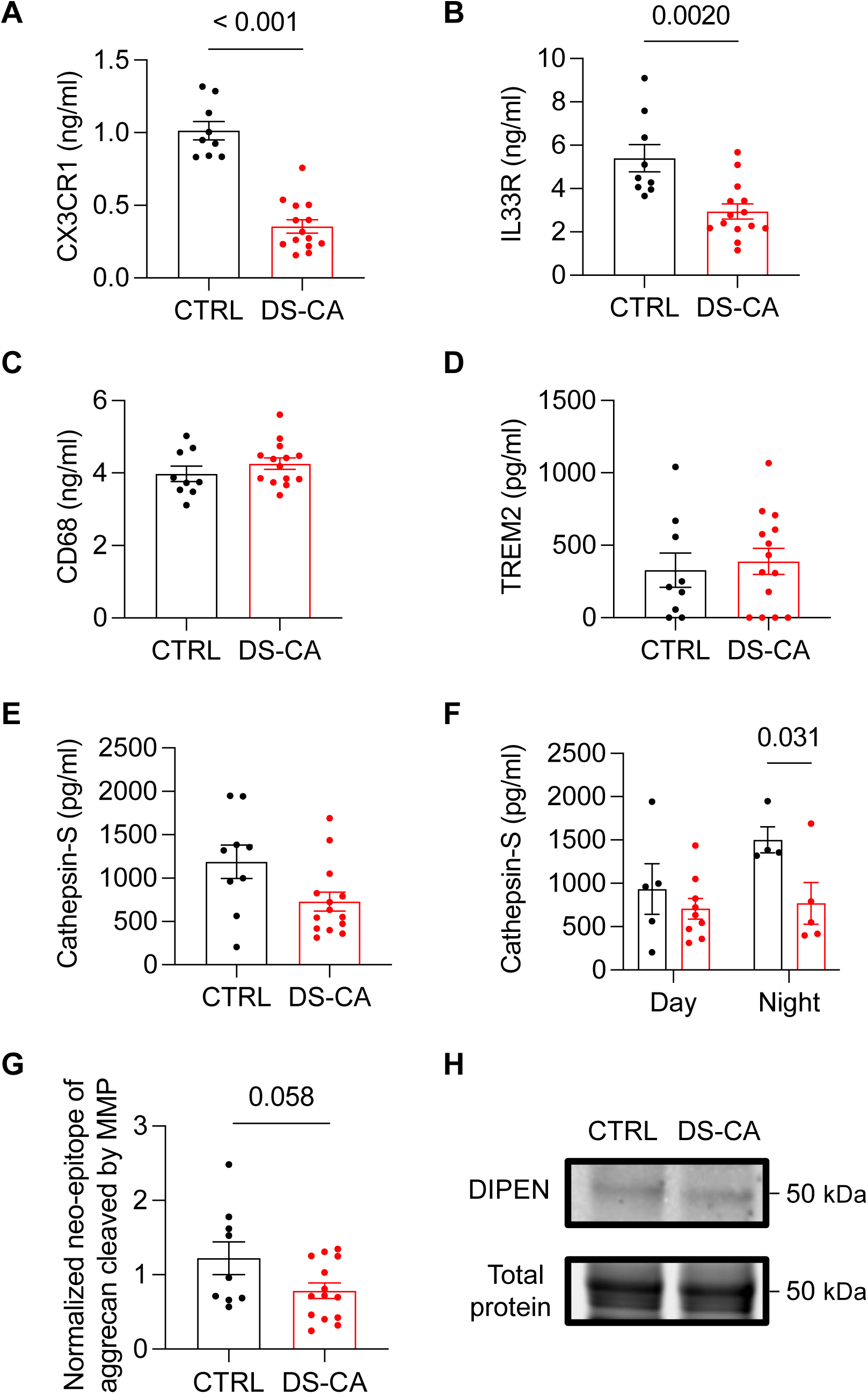
Microglia/macrophage markers involved in PNN regulation are reduced in vmPFC grey matter lysate from depressed suicides with a history of child abuse compared to matched controls. **A** CX3CR1 and **B** IL33R are significantly decreased in DS-CA compared to CTRL as investigated by ELISA. **C** CD68 and **D** TREM2 are not associated with changes associated with child abuse. **E-F** Cathepsin-S levels are significantly lower in DS-CA that died at night (between 6:00PM-5:59AM) **G-H** The cleavage of aggrecan core protein in DS-CA by MMPs at the DIPEN terminal is showing a slight reduction compared to CTRL. CX3CR1: CX3C motif chemokine receptor 1, IL33R: interleukin 33 receptor, CD68: cluster of differentiation 68, TREM2: triggering receptor expressed on myeloid cells 2), ELISA: enzyme-linked immunosorbent assay, MMP: matrix metalloproteinase, CTRL: control, DS-CA: depressed suicide with a history of child abuse.

Lastly, immunoblotting experiments targeting the neo-epitope of aggrecan cleaved by MMPs at the DIPEN terminal (location of cleavage by MMP-2 and MMP-9) in whole grey matter lysate from the vmPFC revealed a slight reduction in the digestion of aggrecan core protein by gelatinases in vmPFC samples from DS-CA individuals (P = 0.058, **Fig. 3G-H**).

### 3.4 Effect of childhood adversity on MMP levels in vmPFC grey matter CD11b^+^ isolated cells

Given the ubiquitous secretion of MMPs and TIMPs from nearly all cell types (Könnecke and Bechmann, 2013), we selectively isolated CD11b^+^ microglia/macrophages with CD11b magnetic beads from fresh brain samples and processed these cells for the MMP array. To estimate the percentage of isolated CD11b^+^ cells that are resident microglia and not infiltrating macrophages after isolation, we investigated the percentage of C11b^+^ cells co-expressing P2RY12, a selective resident microglia marker (Cserép et al., 2020; Zhang et al., 2014), in 10μm-thick sections from the same subjects (CTRL = 4, DS-CA = 4) using fluorescent in situ hybridization (FISH, **Fig. 4A**).

**Figure 4:**
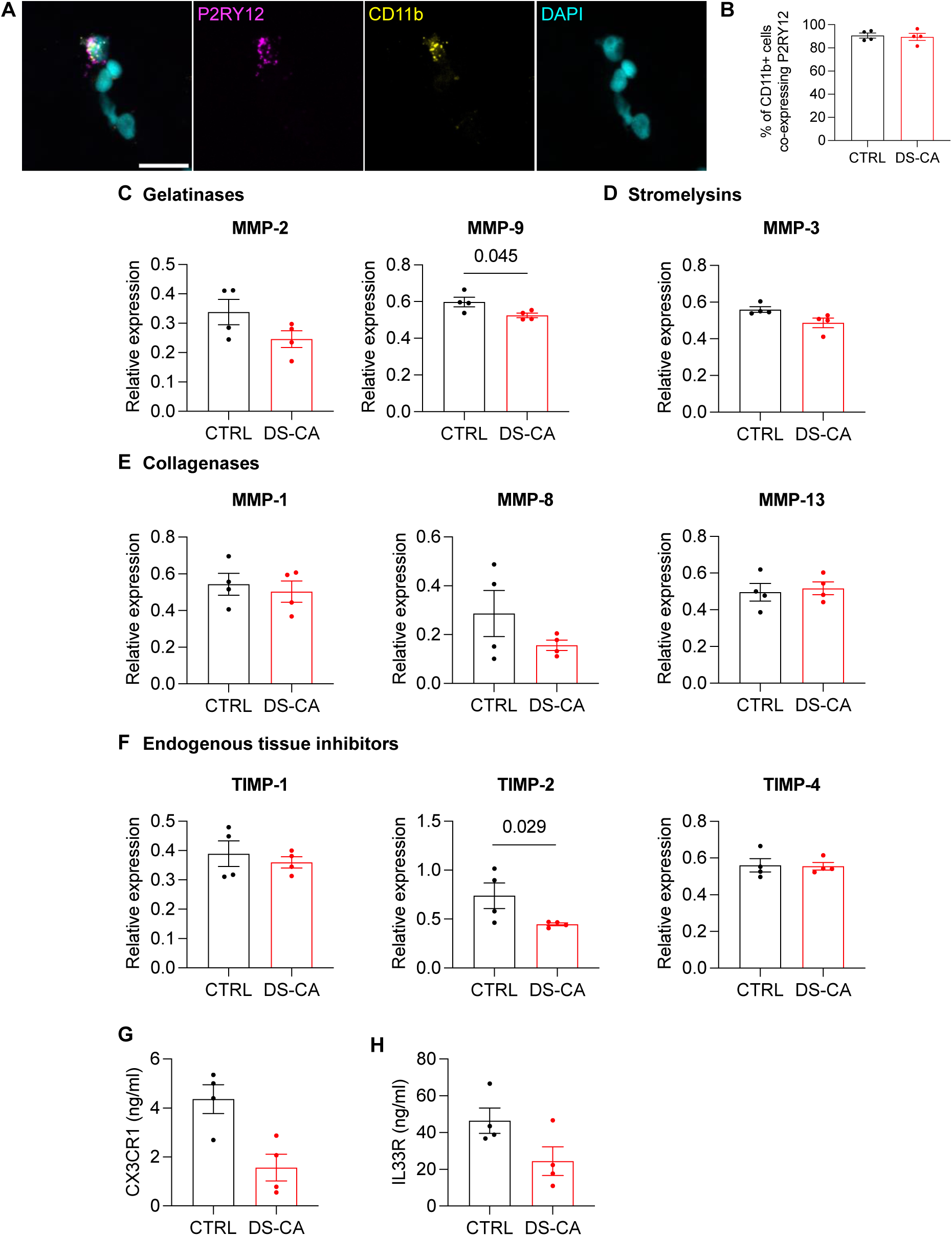
Reduced expression of MMP-9 and TIMP-2 in microglia isolated from vmPFC of depressed suicides with a history of child abuse compared to matched controls. **A** Representative micrograph of fluorescent in situ hybridization for P2RY12 (magenta), CD11b (yellow) and nuclei (cyan) in 10μm-thick sections of vmPFC grey matter. Imaged on an Evident Scientific FV1200 confocal at 40X magnification. Scale bar = 25μm **B** 90% ± 4.9 of CD11b+ cells are P2RY12+ microglia. **C-F** Only MMP-9 and inhibitor TIMP-2 are significantly decreased in microglia isolated from the vmPFC of DS-CA. **G** Microglial CX3CR1 and **H** microglial IL33R are slightly reduced in DS-CA samples compared to CTRL. MMP: matrix metalloproteinase, TIMP: endogenous tissue inhibitor, CTRL: controls, DS-CA: depressed suicide with a history of child abuse.

In all samples, 90% ± 4.9 of CD11b^+^ cells co-expressed P2RY12 (**Fig. 4B**) confirming that our isolated population is indeed enriched for microglial cells. Furthermore, the measurement of MMPs and TIMPs within isolated microglia is plausible as demonstrated by previous studies demonstrating the capacity of microglial translating mRNA to produce these enzymes (Boutej et al., 2017; Rahimian et al., 2024b). In microglial cells isolated from the vmPFC, gelatinase MMP-9 (P = 0.045) and inhibitor TIMP-2 (ANCOVA: F(1,5) = 9.14, P = 0.029) displayed significant downregulation in DS-CA compared to CTRL. Meanwhile, MMP-2 (ANCOVA: F(1,5) = 3.71, P = 0.11), Stromelysin MMP-3 (P = 0.050), all collagenases (MMP-1: P = 0.65, MMP-8: P = 0.27 and MMP-13: P = 0.73), TIMP-1 (P = 0.57) and TIMP-4 (P = 0.91) showed no significant difference between groups (**Fig. 4C-F**). In sum, the concurrent downregulation of MMP-9 and TIMP-2 in both whole grey matter lysate and isolated microglia likely suggest a dysregulation of microglial ECM degradation in DS-CA samples compared to matched controls.

### 3.5 Effect of childhood adversity on the levels of microglia/macrophage markers involved in the regulation of PNNs in vmPFC grey matter CD11b^+^ isolated cells

Seeing as we discovered a significant decrease in CX3CR1 and IL33R in grey matter whole lysate, we next wanted to investigate the same markers in CD11b^+^ isolated microglia. Using ELISA, we found significantly reduced CX3CR1 levels in DS-CA samples (P = 0.01), however, when covariates are included in the model, and with such a small sample size, significance is lost (ANCOVA: F(1,5): 4.14, P = 0.10, **Fig. 4G**). For IL33R, we see a trend in reduced levels in DS-CA (P = 0.08, **Fig. 4H**).

### 3.6 Histological evidence of indirect microglial regulation of PNNs

To better understand the microglia-PNN interaction in our postmortem human brain samples we conducted histological experiments with fresh brain tissue from the same subjects used for CD11b^+^ cell isolation (CTRL =4, DS-CA = 4). To validate our CD68 ELISA results, we immunofluorescently labeled microglia and CD68. Microglia were then categorized based on CD68 high or low expression. We found no difference between groups in expression of CD68 in vmPFC grey matter (F(1,6): 4.3, P = 0.08, **Fig. 5A-B**). Due to the important role of Gas6 in microglial phagocytosis (Grommes et al., 2008) and previous findings indicating a robust dysregulation of Gas6 in the hippocampus of depressed individuals with a history of child abuse which were validated in an animal model of ELS (Reemst et al., 2022), we quantified the percentage of resident microglia (P2RY12^+^ cells) co-expressing Gas6 in vmPFC grey matter using FISH (**Fig. 5C**). We found no difference in the percentage of microglia expressing Gas6 in DS-CA (ANCOVA: F(1, 5) = 0.9, P = 0.39, **Fig. 5D**) compared to controls. To further investigate, we again performed immunofluorescence, labeling microglia, PNNs and parvalbumin-positive inhibitory interneurons in vmPFC. We examined their spatial relationship (**Fig. 5E**) by locating a PNN and tallying the number of microglia found within 50 μm (divided into concentric circles spaced 10 μm apart). We found no difference between groups in number of microglia within a biologically relevant distance from PNNs (F(4,24) = 0.92, P = 0.47, **Fig. 5F**). Taken together the histological and previous ELISA results (CD68 and TREM2) suggest that aberrant phagocytosis is most likely not the mechanism at play.

**Figure 5:**
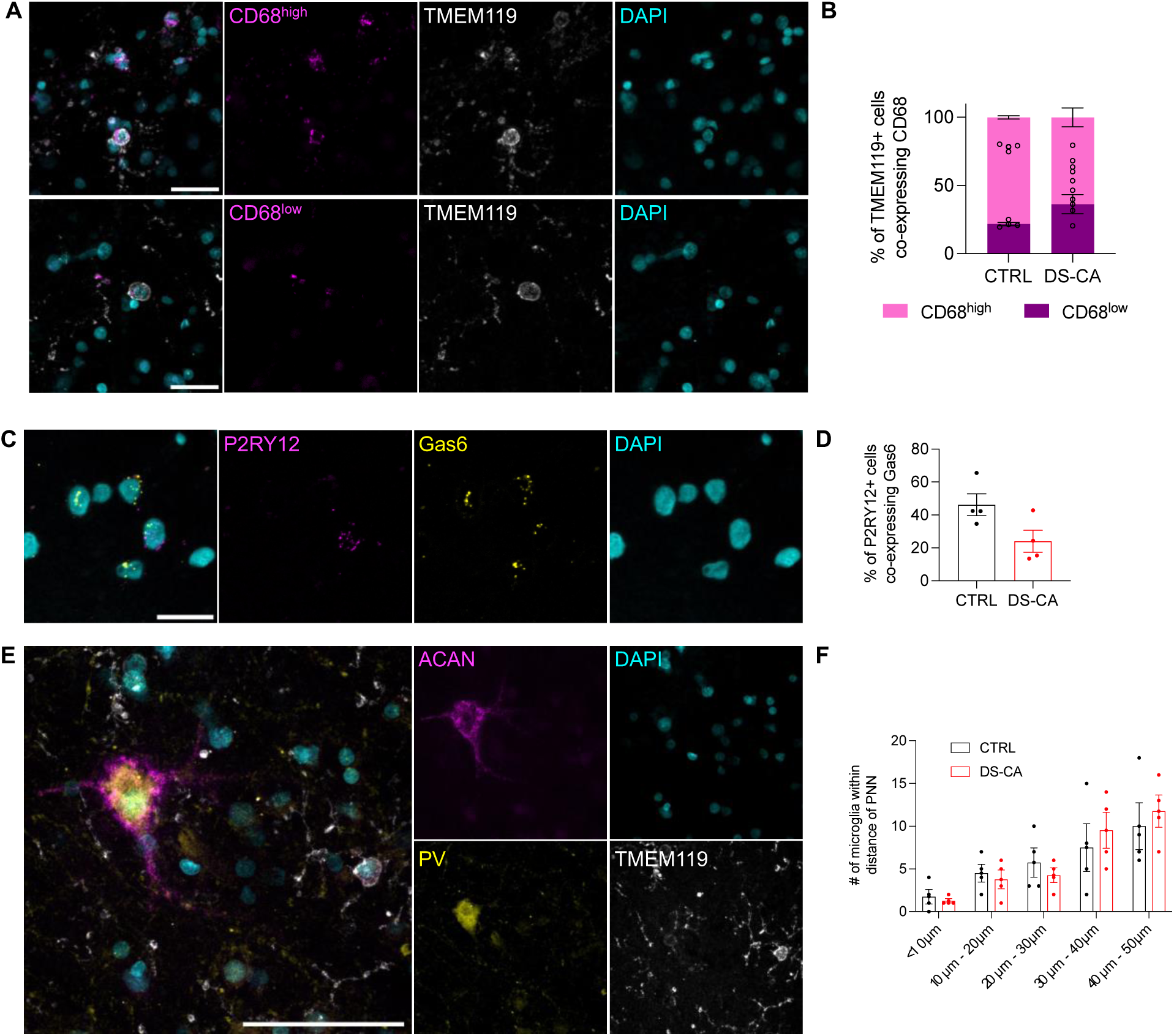
Histological investigation of microglia and PNNs in grey matter vmPFC. **A** Representative micrograph of immunofluorescent labeling of CD68 (magenta), microglia (TMEM119, white) and nuclei (cyan) in fresh-fixed 40μm-thick sections. Imaged on an Evident Scientific FV1200 confocal at 40X. Scale bar = 25μm **B** CD68 expression did not vary between groups. **C** Representative micrograph of fluorescent in situ hybridization for P2RY12 (magenta), Gas6 (yellow) and nuclei (cyan) in 10μm-thick sections of vmPFC grey matter. Imaged on an Evident Scientific FV1200 confocal at 40X magnification. Scale bar = 25μm **D** The percentage of cells expressing Gas6 did not differ between groups **E** Representative micrograph of immunofluorescent labeling of PNNs (aggrecan, magenta), parvalbumin-positive inhibitory neurons (yellow), microglia (TMEM119, white) and nuclei (cyan) in fresh-fixed 40μm-thick sections. Imaged on an Evident Scientific FV1200 confocal at 60X. Scale bar = 50μm **F** Microglia spatial relationship with PNNs is not affected by child abuse. CTRL: controls, DS-CA: depressed suicide with a history of child abuse.

## Discussion

In this study, we sought to explore potential mechanisms underlying the child abuse-associated increase in vmPFC PNNs recently reported by our group (Tanti et al., 2022). We focused mainly on microglial factors, and more specifically on MMP-mediated regulation. Our results indicating a significant downregulation of MMPs, TIMPs and other factors involved in the regulation of ECM in vmPFC samples from DS-CA compared to CTRL are consistent with our previous finding of increased PNN density.

Examining the molecular landscape through the MMP antibody array highlighted a reduction in MMP-9 in DS-CA compared to CTRL, which was evident in both vmPFC whole grey matter lysate and CDllb^+^ isolated microglia. Furthermore, the decrease in aggrecan core protein cleavage by MMPs, which was nearly statistically significantly, suggests not only a decline in abundance but also a possible decrease in enzymatic activity of the MMPs. MMPs are synthesized in a pro-form that remains inactive until proteolytic cleavage. Importantly, this cleavage can be regulated by inflammatory mediators including cytokines, chemokines, free radicals and steroids, or even by other MMPs (Laronha and Caldeira, 2020). We also found reduced TIMPs, specifically microglial TIMP-2, which implies a broader dysregulation of the homeostatic system or a negative feedback loop where decreased MMP activity results in the need for fewer inhibitors. While the current study represents the first postmortem assessment of MMPs (pro and active form) in child abuse victims, polymorphism in the MMP-9 gene has previously been suggested to increase susceptibility to depression, although conflicting reports on MMP-9 levels in depressed patients suggest a complex relationship (Beroun et al., 2019; Li et al., 2022b). While our understanding of the relationship between microglial MMP-9 and PNN in depression is limited, studies in rodent models of neurodegeneration such as TDP-43^Q331K^ and SOD1^G93A^ ALS have reported increased PNN engulfment by MMP-9-positive microglia (Cheung et al., 2024a; Cheung et al., 2024b).

In another set of experiments, we measured the levels of various markers implicated in ECM regulation by microglial cells, although it is important to note that these markers can also be expressed by other cell types. For instance, CX3CR1 can also expressed by oligodendrocyte lineage cells and neural progenitor cells (de Almeida et al., 2023). We observed a strong reduction in CX3CR1 and IL33R in bulk grey matter lysate and reduced levels in CD11b^+^ isolated microglia. Fractalkine signaling (neural CX3CL1 and microglial CX3CR1 interaction) plays a prominent role in complement-independent synaptic elimination (Hoshiko et al., 2012). Interestingly, signaling through CX3CR1 also plays a role in regulating fear behaviours as reported in a mouse model of post-traumatic stress disorder (PTSD) (Schubert et al., 2018). Moreover, CX3CR1 deficiency has been shown to impair neuron–microglia communication under chronic unpredictable stress leading to a lack of responsiveness by microglia (Milior et al., 2016). As mentioned previously CX3CR1 is involved in pruning during a sensitive period and therefore it is necessary for glutamatergic synaptogenesis during development (Hoshiko et al., 2012; Ragozzino et al., 2006). Decreased levels of CX3CR1 during a sensitive period is associated with less glutamatergic AMPA activity and delays the developmental switch in GluN2B to GluN2A NMDA receptors (Andersen, 2022; Szepesi et al., 2018). In a pioneering study by Tansley et al., CX3CR1 knockout (KO) mice subjected to sciatic nerve injury exhibited diminished PNN degradation and reduced lysosomal glycosaminoglycan accumulation within microglia, a stark contrast to the outcomes observed in wild-type mice (Tansley et al., 2022). Taken together our results suggest reduced levels of CX3CR1 in the vmPFC of DS-CA might underlie the increased number of PNNs reported in these subjects (Tanti et al., 2022).

Understanding of neuroglial interaction through CX3CL1/CX3CR1 in brain pathologies such as depression could provide new mechanistic insight into develop therapeutic targets. In the current study our chemokine array and ELISA findings show that the level of CX3CL1 is not altered significantly between CTRL and DS-CA groups.

In a similar vein, IL33 signaling through IL33R is necessary for proper central nervous system formation and functioning (Dwyer et al., 2022). During neurodevelopment, astrocytic release of IL33 plays a crucial role in initiating microglial clearing of excessive synapses (Vainchtein et al., 2018). Conversely, the expression of neuronal IL33 is experience-dependent, leading to microglial engulfment of ECM in the adult hippocampus (Nguyen et al., 2020). In our study, we found a child abuse-associated downregulation of IL33R, in a brain area exhibiting a child abuse-associated increase in PNNs (Tanti et al., 2022). This suggests a diminished signaling cascade between IL33 and IL33R, resulting in attenuated ECM digestion, as observed by Nguyen and colleagues (2020) in neuronal IL33 knock-outs (Nguyen et al., 2020).

Another noteworthy enzyme, Cat-S, has been proposed as an ECM regulator influenced by the circadian rhythm (Pantazopoulos et al., 2020). Cat-S secreted from microglia contributes to the diurnal variation in cortical neuron spine density through proteolytic changes of peri-synaptic ECM (Nakanishi, 2020). Both mouse and human brains exhibit diurnal fluctuations of Cat-S expression in microglia that are antiphase to WFL+ PNN densities, with high Cat-S expression being associated with low WFL+ PNN expression. Interestingly, these rhythms are region-specific, as revealed by human thalamic reticular nucleus expressing more WFL+ PNNs at night and fewer during the day, while the amygdala exhibits an inverse pattern (Pantazopoulos et al., 2020). Furthermore, activated Cat-S incubated with mouse brain sections for just 3 hours reduces WFL+ PNNs, while 24 hours eliminates them completely, highlighting microglial Cat-S specific role in PNN degradation (Pantazopoulos et al., 2020). Interestingly Cat-S can promote the generation of soluble CX3CL1 (the CX3CR1 ligand) (Fonović et al., 2013). The lower expression of Cat-S in individuals with a history of child abuse suggests a reduced availability of CX3CL1 for CX3CR1, indicating that child adversity may be associated with both lower Cat-S expression and diminished availability of CX3CL1 for CX3CR1.

During development, microglia exhibit heightened activity, playing a crucial role in shaping and refining immature neural circuits through the regulation of neurogenesis, synaptogenesis and synaptic pruning (Crapser et al., 2021; Wright-Jin and Gutmann, 2019). Changes in microglial function during this developmental period have been shown to impact these processes, influencing behavior. However, the specific impact of early-life adversity on microglia has remained largely unexplored until recently (Bolton et al., 2022; Dayananda et al., 2023; Johnson and Kaffman, 2018). A recent study by the Korosi lab, in collaboration with our group, has shown an enduring effect of ELS, induced by limited bedding and nesting, on the microglial transcriptome and function under basal and immune-challenged conditions in mice (Reemst et al., 2022). These effects were found to persist for up to 6 months after exposure to the stress paradigm. In an *in vitro* assay, this study further revealed that microglia exposed to ELS exhibit a reduced capacity for synaptosome phagocytosis in comparison to control microglia (Reemst et al., 2022). This observation adds a layer of complexity to our understanding of the prolonged impact of early-life adversity on microglial function and its potential implications for synaptic dynamics in the developing brain.

Using an unbiased and sensitive chemokine array, we quantified the protein expression of CX3CL1 (a selective neural ligand for microglial CX3CR1) and several canonical chemokines including CCL2, CCL3, CCL4 and CCL5 in vmPFC grey matter of DS-CA and matched controls. It is well established that these chemokines have instrumental role in attracting different types of peripheral immune cells such as T cells, monocytes/ macrophages, NK cells and immature dendritic cells, to the site of inflammation (Leighton et al., 2018; Syed et al., 2018). Our analyses did not show any significant differences between cases with child abuse history versus controls. These findings align with more recent sn-RNAseq findings which suggest that microglia in the frontal cortex of individuals who died by suicide display a non-inflammatory or even anti-inflammatory phenotype (Maitra et al., 2023). For instance, it has been shown that in dorsolateral prefrontal cortex, female microglia from depressed suicides—but not male microglia—showed significant differences in gene expression compared to controls. Female subjects exhibited a decrease in both pro- and anti-inflammatory immune signaling pathway related genes. They also showed significant negative enrichment scores for inflammation-related reactome pathway gene sets including INF-gamma signaling, IL-4 and IL-13 signaling, IL-10 signaling, and non-canonical NF-KB pathway (Maitra et al., 2023). In a more recent study, Scheepstra and colleagues performed RNA sequencing on acutely isolated post-mortem microglia form non-suicide depressed patients from occipital cortex and found an immune-suppressed grey matter microglial phenotype but not the white matter (Scheepstra et al., 2023).

Gas6 is an important ligand that is expressed by different cell types and has high selectivity for the tyrosine kinase receptors Tyro3, Axl and Mertk (TAM), their interaction induces microglial phagocytosis (Reemst et al., 2022). Dysregulation of this ligand has been shown previously in the hippocampal microglia of mice following ELS and more importantly in the dentate gyrus microglia of depressed suicide with child abuse is (Reemst et al., 2022). The growth factor Gas6 is abundantly expressed in the brain during development and continues to be present in adulthood, potentially serving as a neurotrophic agent for hippocampal neurons. Additionally, the activation of the TAM signaling pathway by secreted ligands like Gas6 suppresses the sustained and unrestricted inherent immune responses of macrophages and microglia. Specifically, the activation of TAM receptors induces the expression of suppressor proteins of cytokine signaling, which either terminate cytokine receptor-mediated signaling or inhibit the transcriptional activity of nuclear factor kappa B (Reemst et al., 2022). In this study we also looked at the percentage of resident microglia (P2RY12^+^ cells) co-expressing Gas6 in grey matter vmPFC of control individuals compared to DS-CA to try and use it as a proxy for microglial phagocytic activity. Intriguingly, the percentage of P2RY12^+^ microglia expressing Gas6 showed a slight child abuse associated reduction. This decrease, along with no change in CD68 levels or expression in situ implies lower levels of phagocytosis in microglia from DS-CA samples. Recently the first comprehensive transcriptomic analysis of intact microvessels isolated from postmortem vmPFC samples from adult healthy controls and DS-CA samples was reported. The findings of this study again indicate a broad child abuse associated downregulation of immune-related pathways in the neurovascular unit (Wakid et al., 2024). We also used immunofluorescence to explore the spatial relationship between microglia and PNNs in our vmPFC samples. We hypothesized that if microglial phagocytosis differed between cases and controls, this spatial relationship would vary accordingly. However, our findings showed no significant difference in the number of microglia surrounding PNNs, whether they encapsulated PV cells specifically or other cell types.

One fundamental question that remains is whether the integrity of neural extracellular matrix is necessary for homeostatic microglia-mediated synaptic remodeling, and if yes which specific pathway would be affected. The answer was not clear until recently. Cangalaya and colleagues studied the influence of the neural ECM on synaptic remodeling by microglia in the retrosplenial cortex of healthy adult mice (Cangalaya et al., 2024). By injecting chABC and recording with in vivo two-photon microscopy, they showed that chABC intervention increased microglial branching complexity and decreased spine removal rate in physiological condition (Cangalaya et al., 2024). Furthermore, reductions in the extracellular matrix impaired synaptic remodeling after synaptic stress caused by photodamage to individual synaptic components. These changes were linked to dysfunctional complement-mediated synaptic pruning, as indicated by decreased deposition of the complement proteins C1q and C3 at synapses and lower expression of the microglial CR3 receptor (Cangalaya et al., 2024). Future studies examining complement-dependent synaptic remodeling and its relationship with PNN integrity, using both rodent models of ELS and postmortem PFC samples, could provide valuable mechanistic insights into how microglia-PNN interactions influence synaptic remodeling during critical developmental periods.

Our study is not without limitations. First, because of limited availability of matched well-characterized samples, the sample size of this study was relatively modest, particularly for experiments isolating microglia from fresh brain samples and subsequent histological experiments. This maybe have prevented us from detecting group differences and making sex- or age-specific observations. Second, our study could not distinguish between microglial subpopulations, which respond to distinct cell types (Hashimoto et al., 2023; Ngozi and Bolton, 2022). For instance, there are microglial subpopulations that specifically respond to inhibitory neurons expressing both *Gabbr1* and *Gabbr2*, revealing GABA_B_ receptiveness (Favuzzi et al., 2021). Furthermore, inhibitory synapse editing in the visual cortex is mediated by microglial MMP-9 (Hashimoto et al., 2023).

It is noteworthy that most of our knowledge on the importance of microglia during development stems from studies on excitatory neurons, with only recent work having been conducted on microglial interactions with inhibitory neurons, such as PV interneurons (Rahimian et al., 2024a). Given that PNNs predominantly surround parvalbumin-expressing inhibitory interneurons, it raises a compelling question: is there a specific link between these microglial subpopulations and the regulation of PNNs surrounding parvalbumin-expressing inhibitory neurons in the vmPFC? Moreover, the ECM can be categorized into various subtypes, such as PNNs, perisynaptic and perinodal ECM (Chelini et al., 2024; Fawcett et al., 2019). In this study, however, we focused our histological analysis exclusively on PNNs, while our molecular investigations explored microglial factors that have the potential to modulate all ECM subtypes. Further studies are needed to investigate the relationship between microglia and the various forms of ECM, using animal models where more controlled experimentation can be conducted.

Despite these limitations, our study provides the first evidence supporting a role for microglia in the child abuse-associated increase in PNNs in the mature vmPFC. Combined with our previous report implicating oligodendrocyte precursor cells in this phenomenon (Tanti et al., 2022), these results further highlight that early-life adversity has a lasting impact on glial cells, with consequences for the neuroplasticity of brain circuitries implicated in cognition and mood.

## Author contributions

**Claudia Belliveau**: Conceptualization, Methodology, Formal Analysis, Investigation, Writing – Original Draft, Visualization. **Reza Rahimian**: Conceptualization, Methodology, Investigation, Writing Original Draft. **Gohar Fakhfouri**: Investigation, Writing – Review & Editing. **Clémentine Hosdey**: Investigation, Software, Formal Analysis, Writing – Review & Editing. **Sophie Simard**: Software, Formal Analysis, Writing – Review & Editing. **Maria Antonietta Davoli**: Investigation, Project Administration, Writing – Review & Editing. **Dominique Mirault**: Resources, Writing – Review & Editing. **Bruno Giros**: Supervision, Writing – Review & Editing. **Gustavo Turecki**: Resources, Writing – Review & Editing. **Naguib Mechawar**: Conceptualization, Resources, Writing – Original Draft, Supervision, Funding Acquisition.

## Acknowledgements

We are deeply indebted to the next of kin who consented to donating the brains of their loved ones. We would like to thank the Molecular and Cellular Microscopy Platform for help with the imaging.

## Data Availability

All data generated for this manuscript are included herein, requests can be made to NM for access.

