## Supplementary Table 1 for "Evidence of microglial involvement in the childhood abuse-associated increase in perineuronal nets in the ventromedial prefrontal cortex"

| Spearman’s Rho |  | **MMP1** | **MMP2** | **MMP3** | **MMP8** | **MMP9** | **MMP13** | **TIMP1** | **TIMP2** | **TIMP4** |
| --- | --- | --- | --- | --- | --- | --- | --- | --- | --- | --- |
| **Age** | Coef | -0.05 | -0.16 | -0.04 | 0.17 | -0.02 | -0.20 | 0.08 | 0.18 | -0.003 |
|  | Sig. | 0.84 | 0.47 | 0.88 | 0.43 | 0.93 | 0.36 | 0.71 | 0.41 | 0.99 |
| **PMI (h)** | Coef | 0.10 | 0.04 | -0.01 | 0.23 | 0.09 | -0.01 | 0.03 | 0.21 | 0.01 |
|  | Sig. | 0.64 | 0.86 | 0.97 | 0.30 | 0.69 | 0.74 | 0.90 | 0.33 | 0.66 |
| **pH** | Coef | -0.08 | -0.08 | -0.07 | -0.33 | -0.17 | 0.01 | 0.03 | 0.02 | -0.23 |
|  | Sig. | 0.71 | 0.74 | 0.74 | 0.12 | 0.44 | 0.98 | 0.90 | 0.93 | 0.29 |
| **Refrigeration delay (h)** | Coef | 0.21 | 0.10 | 0.04 | 0.35 | 0.17 | 0.05 | 0.16 | 0.33 | 0.16 |
|  | Sig. | 0.33 | 0.66 | 0.86 | 0.10 | 0.44 | 0.82 | 0.47 | 0.13 | 0.48 |
| **Type of antidepressants** | Coef | -0.20 | 0.19 | -0.21 | -0.21 | -0.16 | -0.02 | -0.30 | -0.35 | -0.18 |
|  | Sig. | 0.38 | 0.40 | 0.35 | 0.35 | 0.47 | 0.94 | 0.18 | 0.12 | 0.41 |
| **Number of medications** | Coef | -0.19 | 0.18 | -0.19 | -0.24 | -0.16 | -0.06 | -0.32 | -0.49 | -0.16 |
|  | Sig. | 0.41 | 0.41 | 0.39 | 0.28 | 0.47 | 0.81 | 0.15 | 0.021 | 0.49 |

Correlation coefficients and P values for Spearman’s rho non-parametric measure of association between co-variates and dependent variables in archived frozen brain samples. Red = correlation is significant at the 0.05 level (2-tailed)

| Spearman’s Rho |  | **CX3CR1** | **CX3CL1** | **IL33R** | **CD68** | **TREM2** | **Cat-S** | **Cleaved aggrecan** | **CCL2** | **CCL3** | **CCL4** | **CCL5** |
| --- | --- | --- | --- | --- | --- | --- | --- | --- | --- | --- | --- | --- |
| **Age** | Coef | 0.02 | -0.34 | 0.35 | -0.37 | -0.36 | -0.12 | 0.03 | -0.18 | -0.03 | -0.10 | -0.12 |
|  | Sig. | 0.94 | 0.11 | 0.10 | 0.08 | 0.09 | 0.59 | 0.91 | 0.42 | 0.88 | 0.66 | 0.60 |
| **PMI (h)** | Coef | -0.02 | 0.05 | 0.15 | 0.23 | -0.20 | 0.16 | 0.15 | -0.06 | -0.003 | 0.04 | 0.07 |
|  | Sig. | 0.92 | 0.82 | 0.49 | 0.29 | 0.36 | 0.47 | 0.49 | 0.77 | 0.99 | 0.88 | 0.75 |
| **pH** | Coef | -0.08 | 0.48 | -0.52 | 0.26 | 0.78 | -0.19 | -0.27 | 0.34 | 0.30 | 0.16 | 0.47 |
|  | Sig. | 0.72 | 0.03 | 0.01 | 0.23 | < 0.001 | 0.38 | 0.21 | 0.12 | 0.17 | 0.49 | 0.03 |
| **Refrigeration delay (h)** | Coef | 0.01 | -0.06 | 0.23 | 0.03 | -0.10 | 0.07 | 0.02 | 0.19 | 0.006 | 0.06 | 0.02 |
|  | Sig. | 0.95 | 0.77 | 0.29 | 0.90 | 0.65 | 0.76 | 0.94 | 0.38 | 0.980 | 0.77 | 0.94 |
| **Type of antidepressants** | Coef | -0.65 | 0.05 | -0.28 | 0.008 | 0.037 | -0.35 | -0.33 | 0.34 | 0.20 | 0.24 | 0.05 |
|  | Sig. | < 0.001 | 0.82 | 0.20 | 0.97 | 0.87 | 0.12 | 0.13 | 0.12 | 0.38 | 0.28 | 0.81 |
| **Number of medications** | Coef | -0.59 | -0.05 | -0.31 | -0.15 | -0.13 | -0.29 | -0.41 | 0.30 | 0.13 | 0.15 | -0.06 |
|  | Sig. | 0.004 | 0.81 | 0.16 | 0.49 | 0.55 | 0.19 | 0.06 | 0.18 | 0.57 | 0.50 | 0.78 |
