## Supplementary Table 2 for "Evidence of microglial involvement in the childhood abuse-associated increase in perineuronal nets in the ventromedial prefrontal cortex"

| Spearman’s Rho |  | **MMP1** | **MMP2** | **MMP3** | **MMP8** | **MMP9** | **MMP13** | **TIMP1** | **TIMP2** | **TIMP4** | **CX3CR1** | **IL33R** |
| --- | --- | --- | --- | --- | --- | --- | --- | --- | --- | --- | --- | --- |
| **Age** | Coef | -0.14 | -0.20 | -0.08 | 0.60 | -0.04 | 0.07 | -0.07 | -0.22 | -0.06 | 0.29 | 0.30 |
|  | Sig. | 0.73 | 0.63 | 0.84 | 0.12 | 0.93 | 0.87 | 0.87 | 0.61 | 0.89 | 0.49 | 0.47 |
| **PMI (h)** | Coef | 0.29 | 0.76 | 0.48 | 0.36 | 0.50 | 0.62 | 0.67 | 0.76 | -.05 | -0.24 | 0.10 |
|  | Sig. | 0.49 | 0.03 | 0.23 | 0.39 | 0.21 | 0.10 | 0.07 | 0.03 | 0.91 | 0.57 | 0.82 |
| **pH** | Coef | 0.61 | -0.14 | 0.07 | -0.14 | 0.36 | -0.04 | 0.21 | 0.14 | 0.54 | 0.32 | 0.00 |
|  | Sig. | 0.15 | 0.76 | 0.88 | 0.76 | 0.43 | 0.94 | 0.65 | 0.76 | 0.22 | 0.48 | 1.00 |
| **Refrigeration delay (h)** | Coef | 0.12 | 0.38 | 0.05 | 0.24 | 0.00 | 0.31 | -0.17 | 0.17 | -0.05 | -0.07 | 0.00 |
|  | Sig. | 0.78 | 0.35 | 0.91 | 0.57 | 1.00 | 0.46 | 0.69 | 0.69 | 0.91 | 0.87 | 1.00 |
| **Type of antidepressants** | Coef | 0.11 | -0.18 | -0.65 | -0.03 | -0.44 | 0.21 | 0.21 | -0.26 | 0.07 | -0.83 | -0.39 |
|  | Sig. | 0.80 | 0.67 | 0.08 | 0.95 | 0.28 | 0.62 | 0.62 | 0.53 | 0.87 | 0.01 | 0.35 |
| **Number of medications** | Coef | -0.04 | -0.13 | -0.54 | 0.08 | -0.58 | 0.24 | 0.24 | -0.26 | -0.11 | -0.85 | -0.55 |
|  | Sig. | 0.93 | 0.76 | 0.17 | 0.85 | 0.14 | 0.58 | 0.58 | 0.53 | 0.81 | 0.008 | 0.16 |

Correlation coefficients and P values for Spearman’s rho non-parametric measure of association between co-variates and dependent variables in fresh brain samples used for microglia isolation. Red = correlation is significant at the 0.05 level (2-tailed)
