## Supplementary Table 3 for "Evidence of microglial involvement in the childhood abuse-associated increase in perineuronal nets in the ventromedial prefrontal cortex"

| Spearman’s Rho |  | **%CD68TMEM119** | **%P2RY12_gas6** | **%CD11b_ P2RY12** |
| --- | --- | --- | --- | --- |
| **Age** | Coef | -0.08 | 0.12 | 0.30 |
|  | Sig. | 0.84 | 0.79 | 0.47 |
| **PMI (h)** | Coef | 0.38 | -0.05 | -0.29 |
|  | Sig. | 0.35 | 0.91 | 0.49 |
| **pH** | Coef | -0.43 | -0.50 | 0.46 |
|  | Sig. | 0.34 | 0.25 | 0.29 |
| **Refrigeration delay (h)** | Coef | 0.31 | -0.33 | -0.67 |
|  | Sig. | 0.46 | 0.42 | 0.07 |
| **Type of antidepressants** | Coef | -0.60 | -0.83 | 0.04 |
|  | Sig. | 0.12 | 0.01 | 0.92 |
| **Number of medications** | Coef | -0.53 | -0.45 | -0.28 |
|  | Sig. | 0.187 | 0.26 | 0.51 |

Correlation coefficients and P values for Spearman’s rho non-parametric measure of association between co-variates and dependent variables in fresh brain samples used for microglia isolation. Red = correlation is significant at the 0.05 level (2-tailed)
