## Supplementary figures and images for "Evidence of microglial involvement in the childhood abuse-associated increase in perineuronal nets in the ventromedial prefrontal cortex"

### Supplementary File 1

Group A

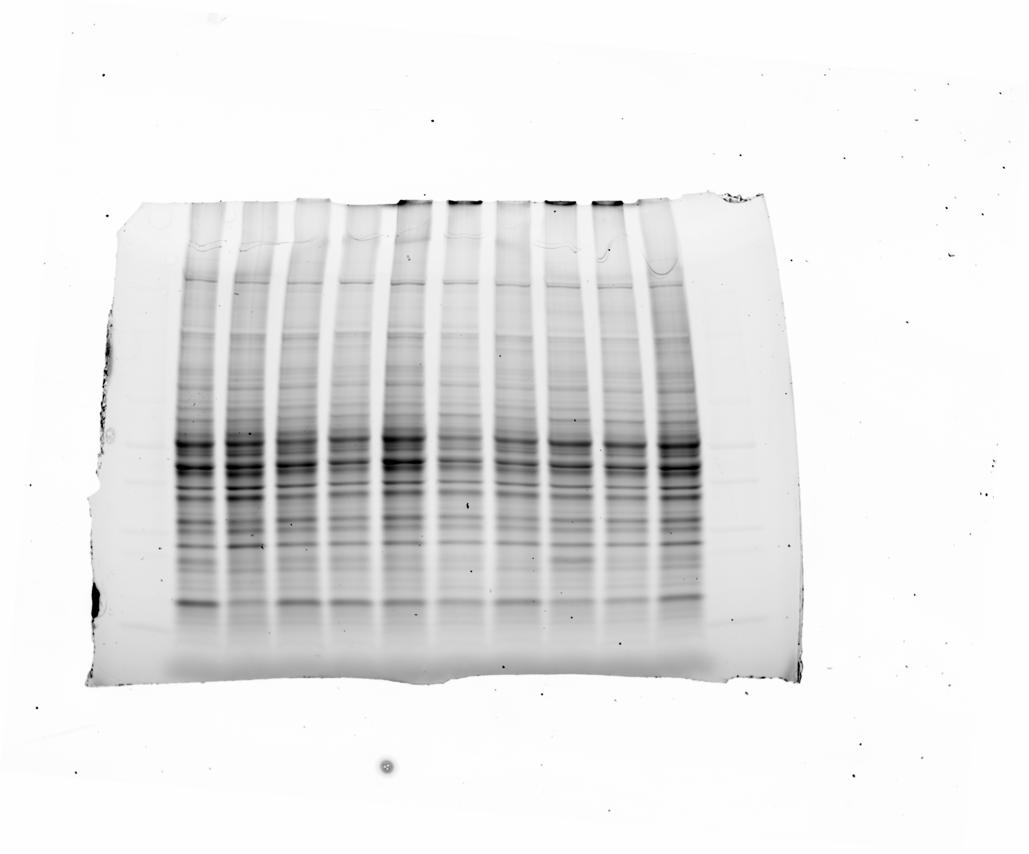

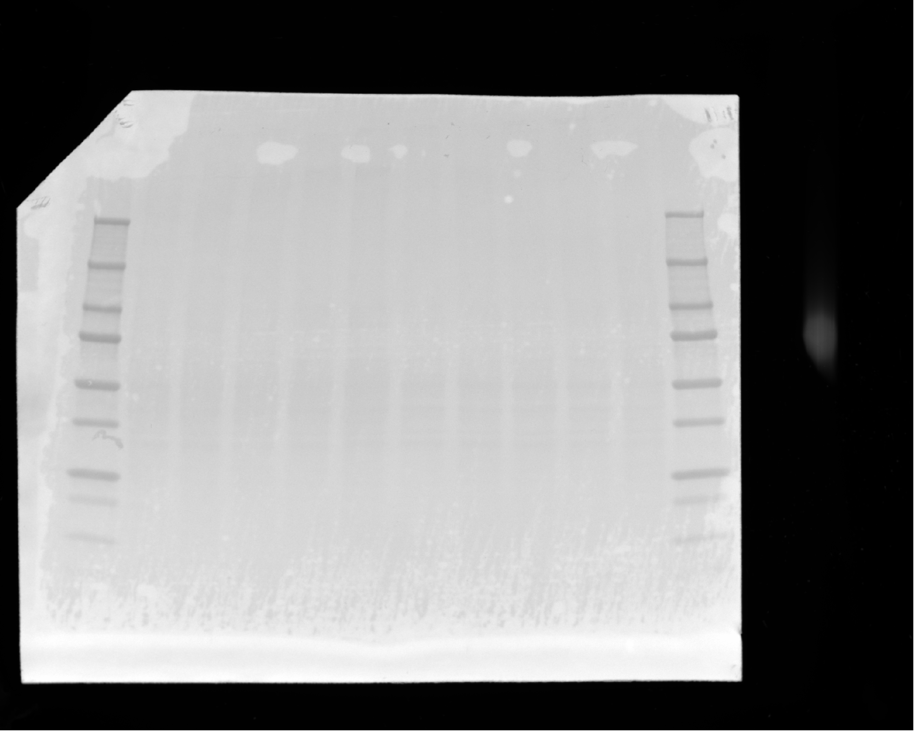

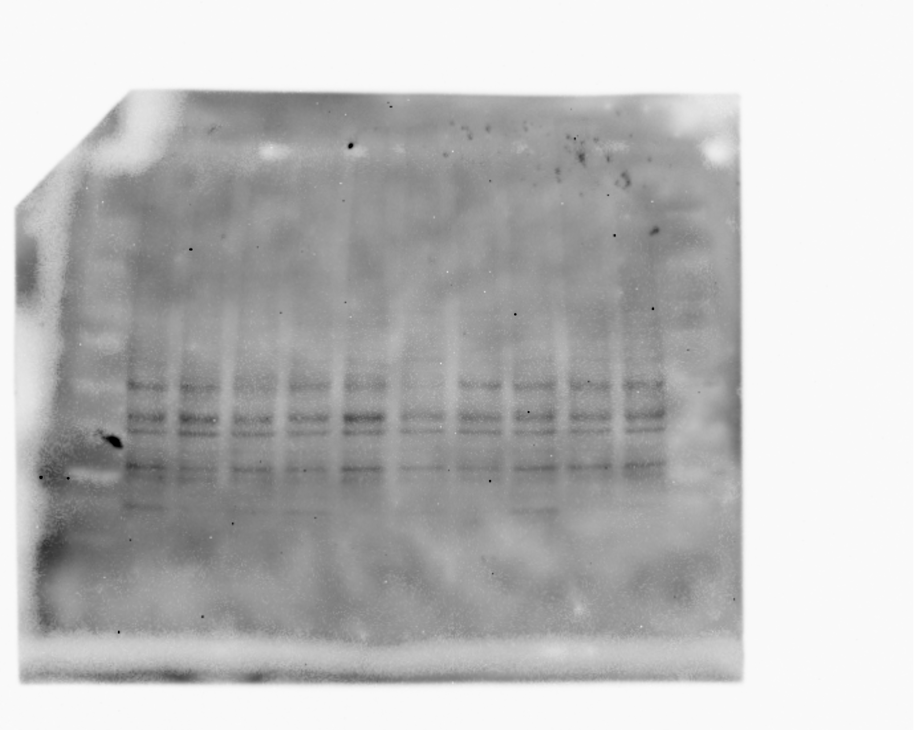

Group B

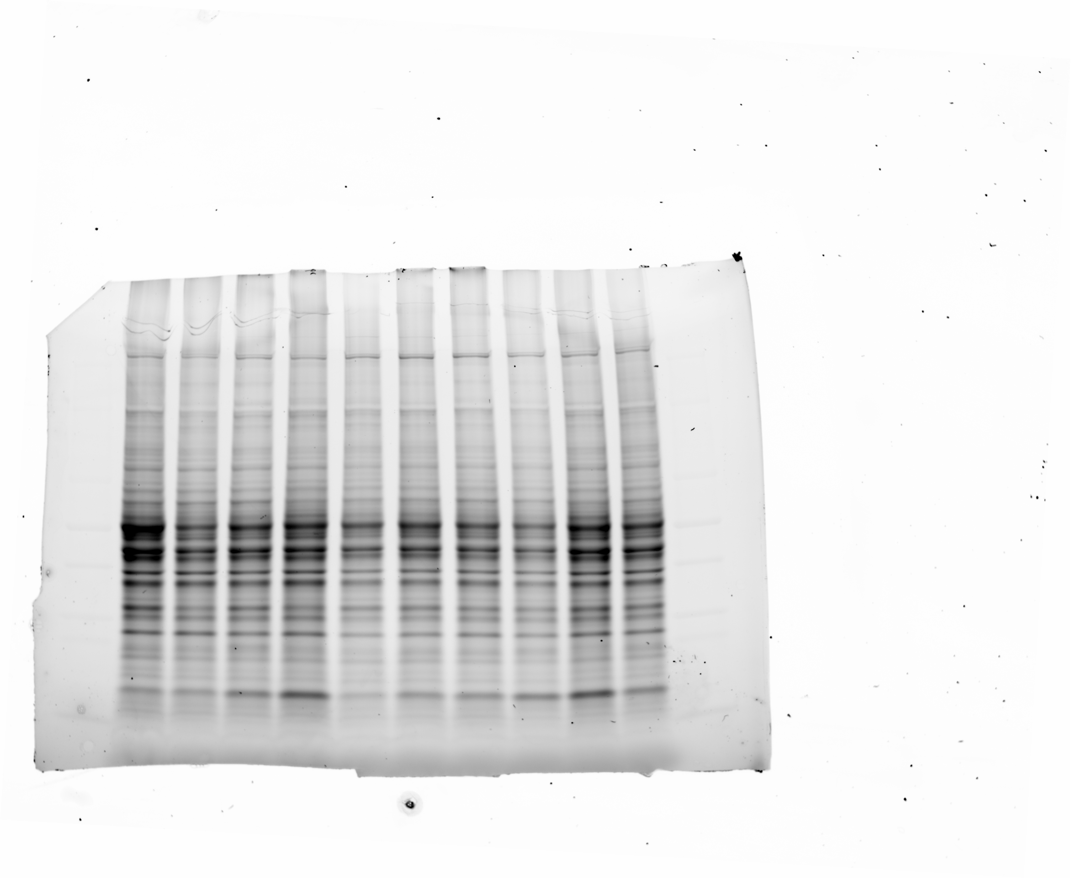

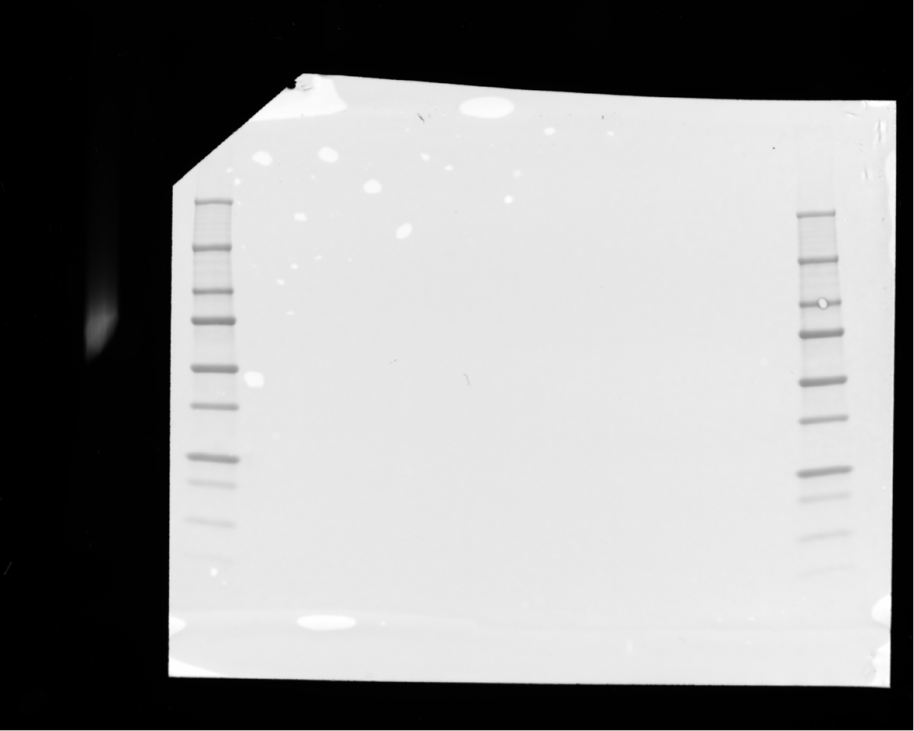

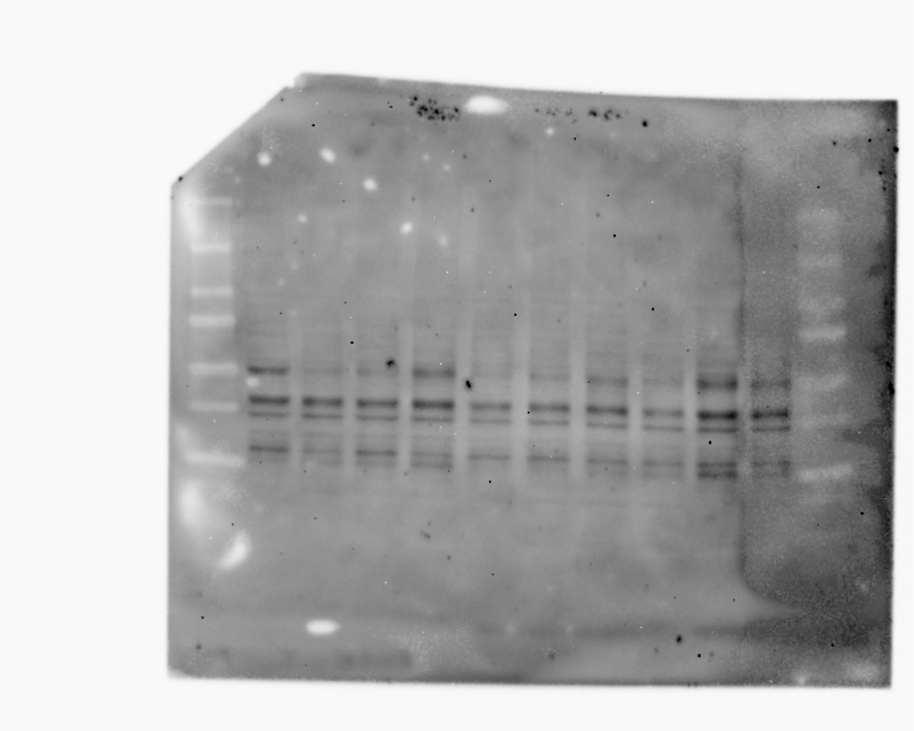

Group C

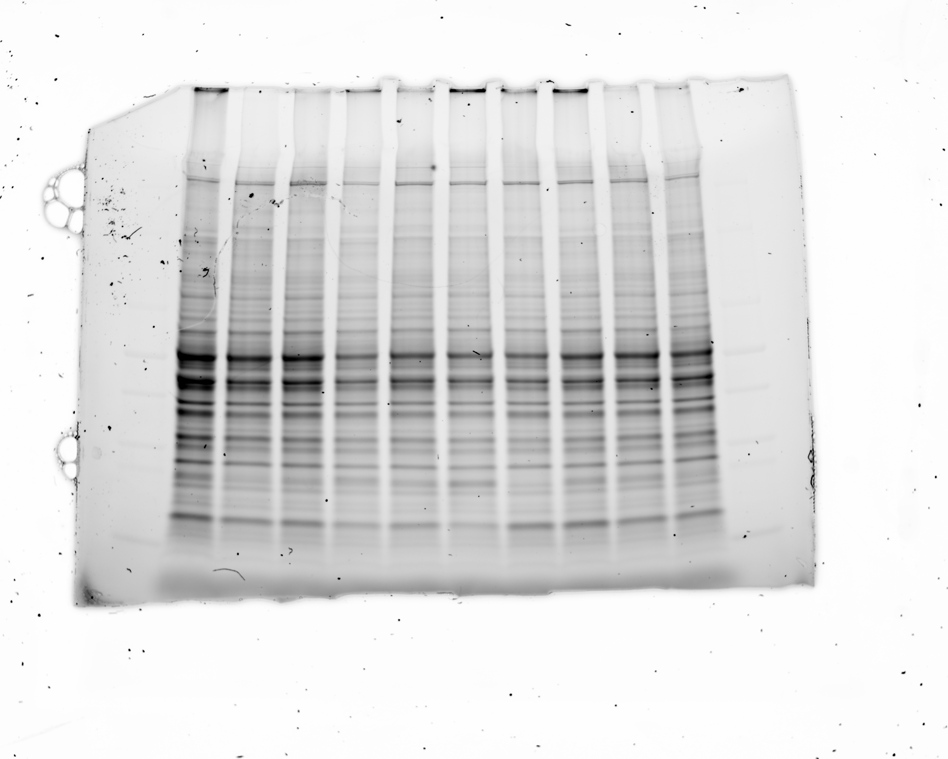

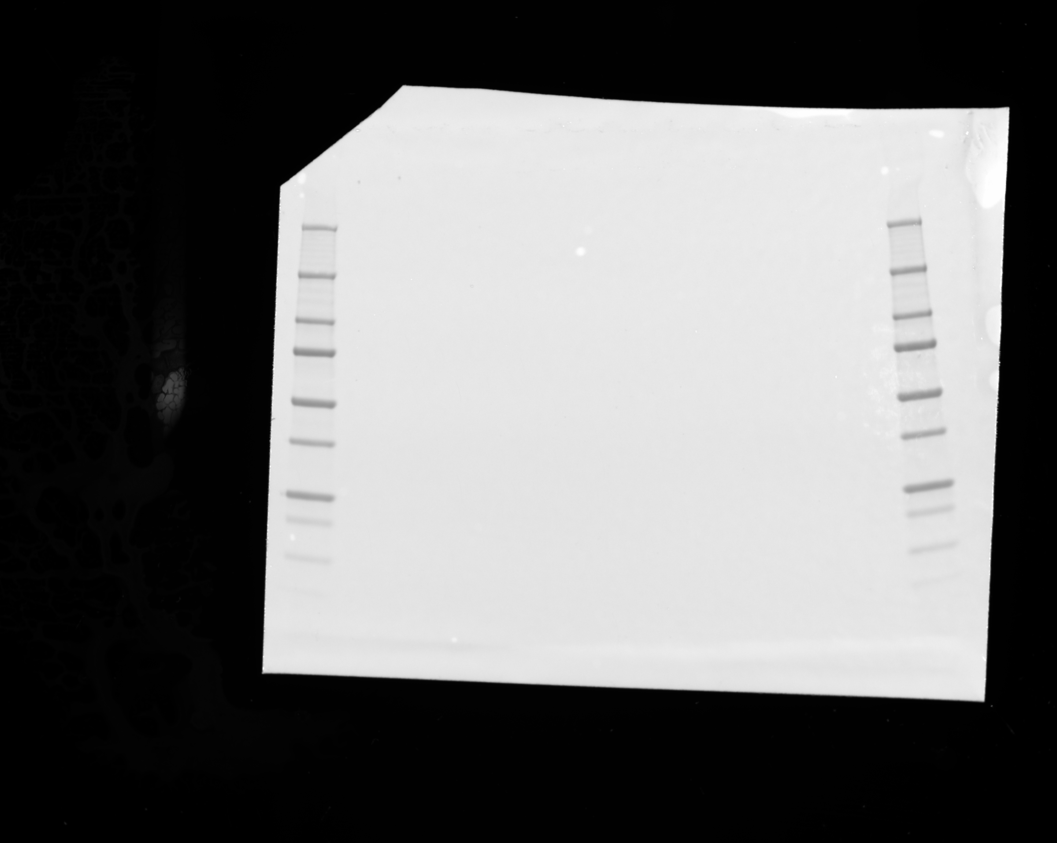

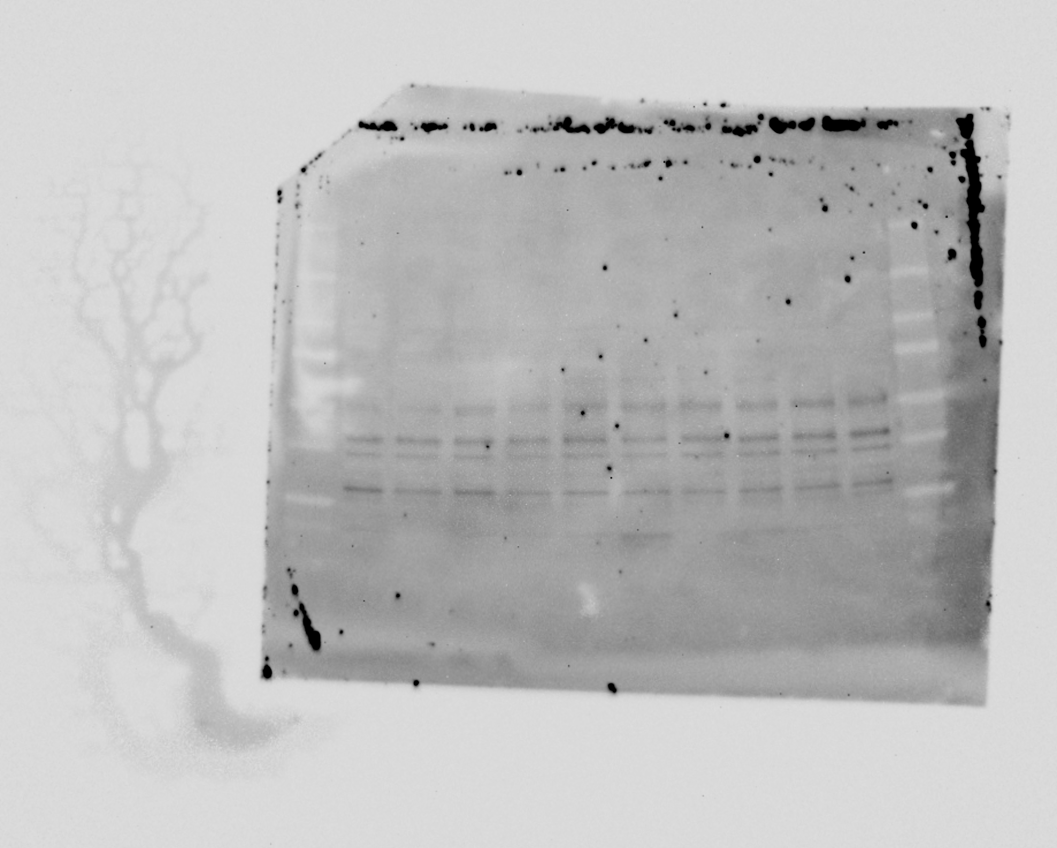

Group D

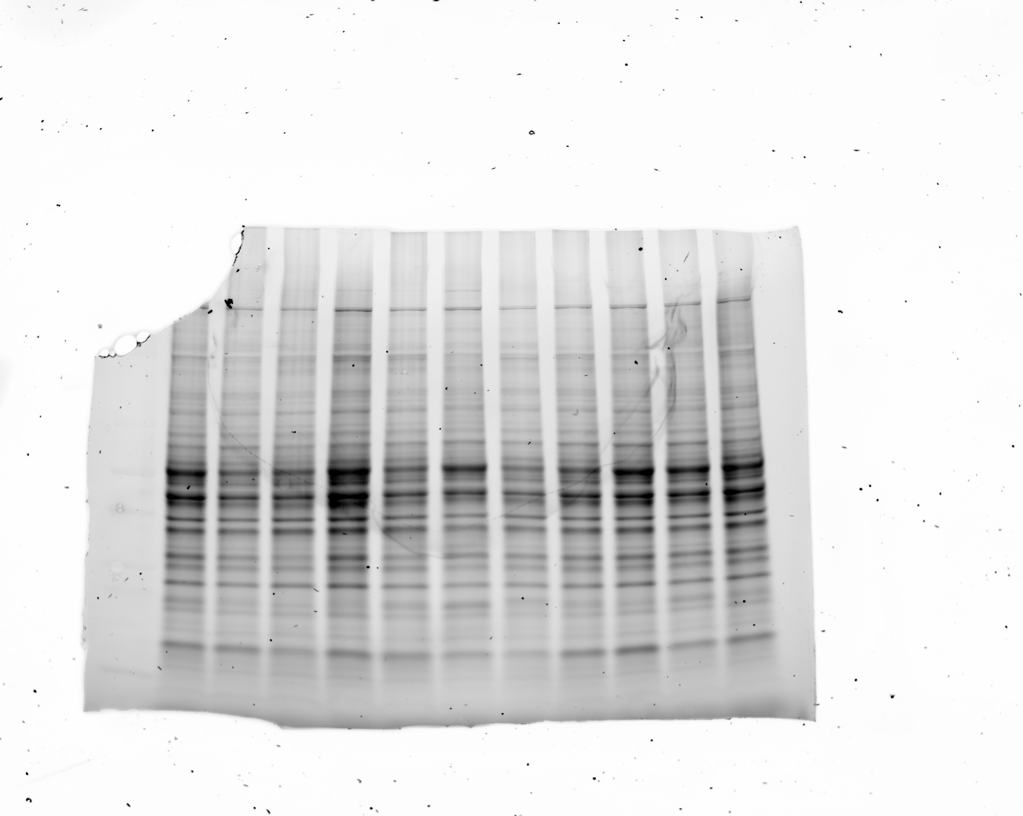

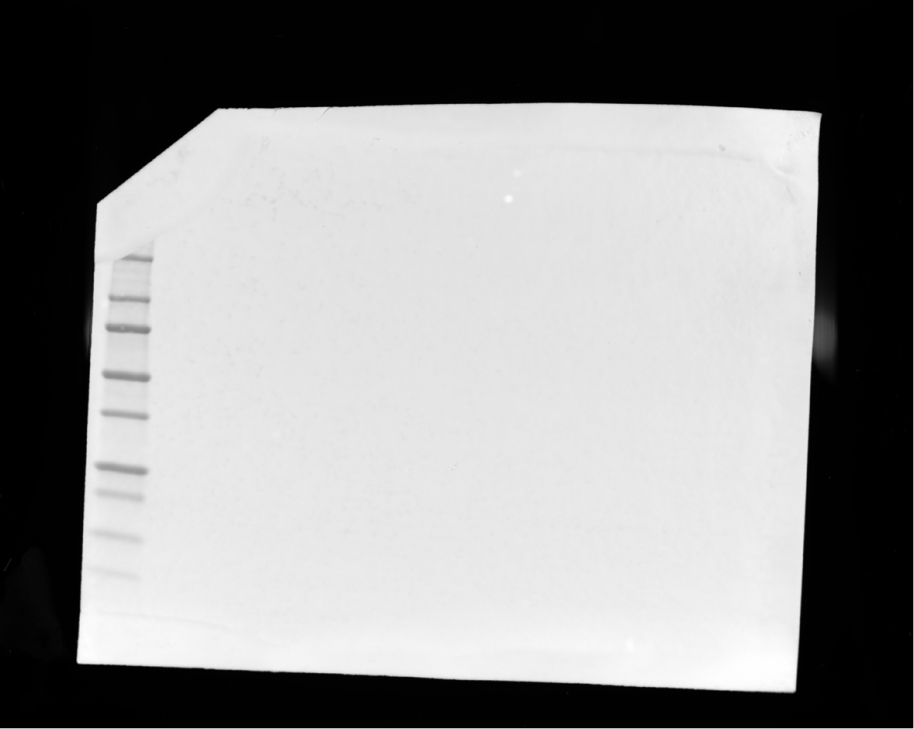

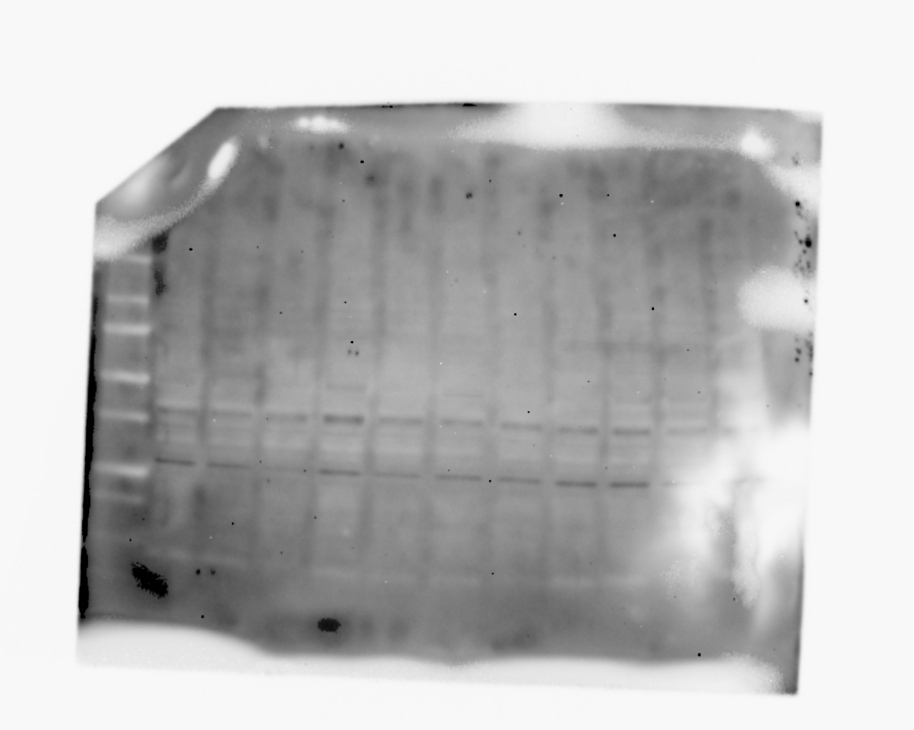

Representative for figure (Groups A,B,C,D pooled by group)

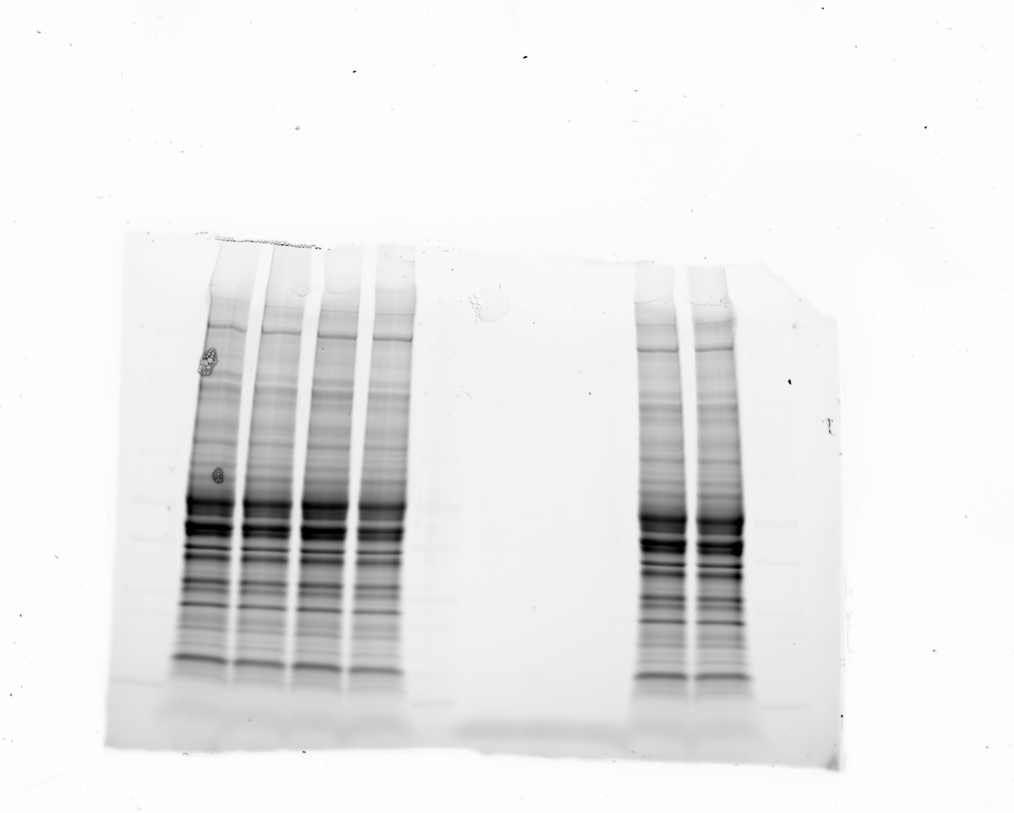

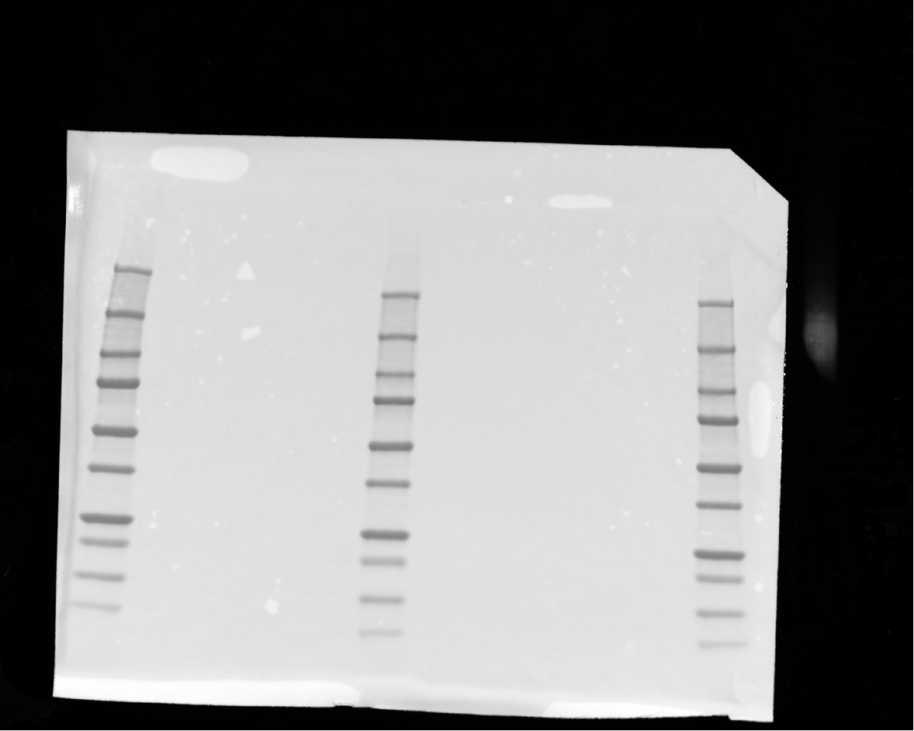

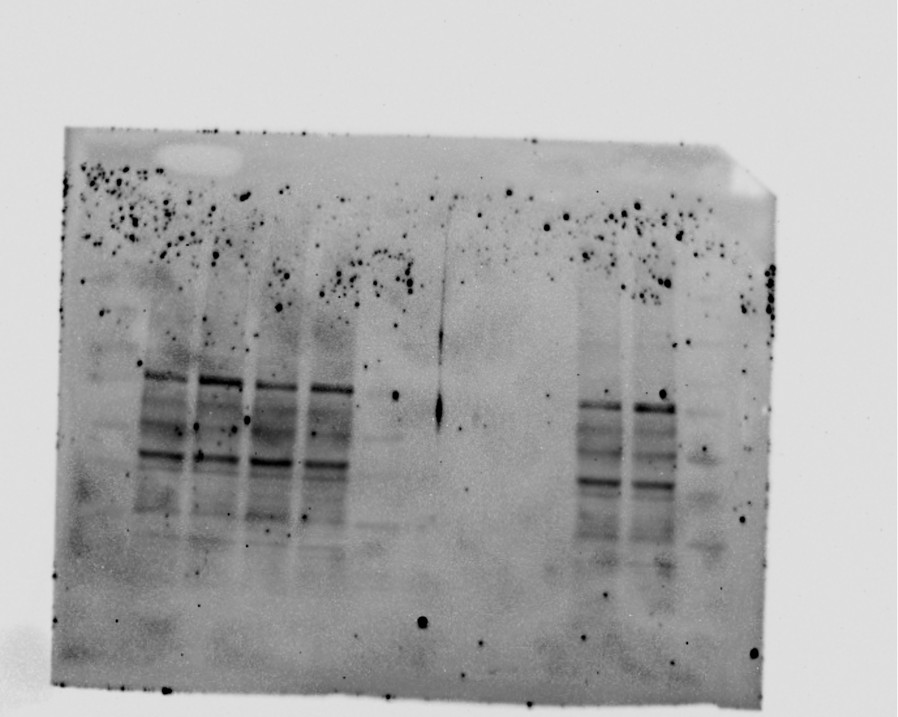
