## Supplementary File 2 for "Evidence of microglial involvement in the childhood abuse-associated increase in perineuronal nets in the ventromedial prefrontal cortex": cell.py

```
from enum import Enum

class Cell:
    def __init__(self, id, position_x, position_y, type, sheetname):
        self.id = id
        self.position_x = position_x
        self.position_y = position_y
        self.type = type
        self.sheetname = sheetname
```
