## Supplementary File 2 for "Evidence of microglial involvement in the childhood abuse-associated increase in perineuronal nets in the ventromedial prefrontal cortex": generate_distances.py

```
from math import dist
import openpyxl
from cell import Cell
import pandas as pd 


# Constants
PATH_DATA = r"/data/PNN_Microglia_Coordinates.xlsx"
PATH_RESULTS = r"/data/PNN_Microglia_results.xlsx"
MAX_DISTANCE = 50 # Distance threshold for analysis (in micrometers, can be adjusted based on interest)
MAX_COLUMN_NUMBER = 30 # Magic number for columns
MAX_ROW_NUMBER = 2000 # Magic number for rows

# Load workbooks
wb_data = openpyxl.load_workbook(PATH_DATA)
wb_results = openpyxl.load_workbook(PATH_RESULTS)

# Intialize dictionaries for subject-specific and region-specific data
sorted_cells = {}

def initialize_dict_structure(roi: str, subject: str, cell_type: str):
    if roi not in sorted_cells:
        sorted_cells[roi] = {}
    if subject not in sorted_cells[roi]:
        sorted_cells[roi][subject] = {}
    if cell_type not in sorted_cells[roi][subject]:
        sorted_cells[roi][subject][cell_type] = []
        
def add_cell_to_sorted_cells(roi: str, subject: str, cell: Cell):
    cell_type = cell.type.lower()
    initialize_dict_structure(roi, subject, cell_type)
    
    if cell_type == "microglia":
        index = len(sorted_cells[roi][subject][cell_type]) + 1
        cell.id = f"Microglia - {roi}#{index}"
    elif cell_type in ["pnnpv", "pnnother"]:
        index = len(sorted_cells[roi][subject][cell_type]) + 1
        cell.id = f"{cell_type.upper()} - {roi}#{index}"
    
    sorted_cells[roi][subject][cell_type].append({
        'id': cell.id,
        'x': cell.position_x,
        'y': cell.position_y
    })
    
def process_sheets():
    for sheet in wb_data.worksheets:
        subject_name = sheet.title
        wb_data.active = sheet
        row_count = sheet.max_row
        print(f"Processing sheet: {subject_name}")
        
        for row in range(2, row_count + 1):
            curr_name = sheet.cell(row=row, column=1).value
            curr_x = sheet.cell(row=row, column=2).value
            curr_y = sheet.cell(row=row, column=3).value
            
            if not curr_name or curr_x is None or curr_y is None:
                continue
            
            curr_roi = curr_name.split('-')[-1].strip()
            curr_type = curr_name.split('-')[0].strip()
            
            cell = Cell(curr_name, curr_x, curr_y, curr_type, subject_name)
            
            add_cell_to_sorted_cells(curr_roi, subject_name, cell)

# Function to generate distances and save results to excel
def generate_distances():

    combinations = {
        'ROI1': {
            'pnnpv-microglia': 'Distance ROI1 PNNPV',
            'pnnother-microglia': 'Distance ROI1 PNNother'
        },
        'ROI2': {
            'pnnpv-microglia': 'Distance ROI2 PNNPV',
            'pnnother-microglia': 'Distance ROI2 PNNother'
        }
    }
    
    for roi in combinations:
        for combination in combinations[roi]:
            sheet_name = combinations[roi][combination]
            if sheet_name not in wb_results.sheetnames:
                wb_results.create_sheet(title=sheet_name)
            # Initialize the sheet with headers
            results_sheet = wb_results[sheet_name]
            results_sheet.delete_cols(1, MAX_COLUMN_NUMBER)
            results_sheet.delete_rows(1, MAX_ROW_NUMBER)
            results_sheet.cell(row=1, column=1).value = "PNN-type and ID"
            results_sheet.cell(row=1, column=2).value = "Microglia ID"
            results_sheet.cell(row=1, column=3).value = "Distance (µm)"
            results_sheet.cell(row=1, column=4).value = "Subject"

    # Calculate distances and write to corresponding sheets
    for roi, subjects in sorted_cells.items():
        for subject, cell_types in subjects.items():
            for (pnn_type, microglia_type) in [('pnnpv', 'microglia'), ('pnnother', 'microglia')]:
                if pnn_type in cell_types and microglia_type in cell_types:
                    # Get the sheet based on ROI and combination
                    sheet_name = combinations[roi][f"{pnn_type}-{microglia_type}"]
                    results_sheet = wb_results[sheet_name]
                    row = results_sheet.max_row + 1
                    for pnn_cell in cell_types[pnn_type]:
                        for microglia_cell in cell_types[microglia_type]:
                            distance = dist(
                                (pnn_cell['x'], pnn_cell['y']),
                                (microglia_cell['x'], microglia_cell['y'])
                            )
                            if distance <= MAX_DISTANCE:
                                results_sheet.cell(row=row, column=1).value = pnn_cell['id']
                                results_sheet.cell(row=row, column=2).value = microglia_cell['id']
                                results_sheet.cell(row=row, column=3).value = distance
                                results_sheet.cell(row=row, column=4).value = subject
                                row += 1

    # Save the results to the Excel file
    wb_results.save(PATH_RESULTS)
    print(f"Distances saved in results sheet: {results_sheet.title}")

## Call the funcitons   
process_sheets()
generate_distances()
print("Distances have been generated in Excel Sheet")
```
